# Lifespan can be extended during a specific time window early in life

**DOI:** 10.1101/2022.02.18.480980

**Authors:** G. Aiello, C. Sabino, D. Pernici, M. Audano, F. Antonica, M. Gianesello, A. Quattrone, N. Mitro, A. Romanel, A. Soldano, L. Tiberi

## Abstract

Lifespan is determined by complex and tangled mechanisms that are largely unknown. The early postnatal stage has been proposed to play a role in lifespan, but its contribution is still controversial. Here, we found that a short rapamycin treatment during early life can prolong lifespan in *Mus musculus* and *Drosophila melanogaster*. Notably, the same treatment at later time points has no evident effect on lifespan, suggesting that we found a crucial time-window involved in lifespan modulation. We discovered that sulfotransferases are upregulated during early rapamycin treatment both in newborn mice and *Drosophila* larvae. Furthermore, overexpression of the sulfotransferase *dST1* triggers an increment in the lifespan of *Drosophila melanogaster*. Our findings unveil a novel link between early-life treatments and long-term effects on lifespan.

**One Sentence Summary:** Early life events increase lifespan.

## Main Text

Genetic and environmental conditions in early organism development could influence traits later in life, including diseases, ageing and lifespan (1–3). Indeed, few studies have reported that different diets for pregnant mothers or for young mice affect offspring survival (3, 4). Notably, there is also controversial evidence about the correlation between early-life treatments and lifespan extension in mammals (3, 5). Similar experiments conducted in *Drosophila melanogaster* and *Caenorhabditis elegans* showed that dietary or changes in cellular ROS levels during development can induce lifespan extension (6–8). Interestingly, lifespan can be also modulated through the regulation of the mTOR pathway. This signaling is evolutionarily conserved from yeast to mammals and regulates growth and metabolism in response to growth factors, amino acids, stresses, as well as changes in cellular energy status (9). Inhibition of the mTOR signaling pathway by genetic or pharmacological intervention extends lifespan in vertebrates, yeast, nematodes, and fruit flies (10–12). Treating mice with rapamycin, an inhibitor of the mTOR pathway, from 20 months of age extends the median and maximal lifespan of both male and female mice (13) and the same effect has been observed in *Drosophila melanogaster* (14). Nevertheless, most of the published data on dietary interventions and drug treatments have been performed during the adult life of several organisms (10), while early-life rapamycin administration has never been tested in wild type mice (15). Indeed, transient rapamycin administration was only used in elder mice (16, 17) and the mechanisms behind this lifespan increase are elusive. Here, we investigated whether an early-life and transient rapamycin administration can prolong lifespan in two animal models, *Mus musculus* and *Drosophila melanogaster*.

## Results

We tested whether mice lifespan was sensitive to early-life modulation by performing an early transient rapamycin treatment on CD1 outbred mice. Rapamycin (10mg/kg) was administrated daily in two distinct temporal windows, from postnatal day 4 to postnatal day 30 (P4-P30), or from postnatal day 30 to postnatal day 60 (P30-P60) (**Fig. 1A**) and the lifespan was evaluated using a Kaplan-Meier survival curve (log-rank test). Combined data from both sexes showed a 9.6% increase in median lifespan in P4-P30 rapamycin-treated mice compared to control mice (treated with ethanol) and 9.1% increase compared to P30-P60 rapamycin-treated mice (**Fig. 1B, E**). The analysis of each gender separately showed similar result, leading to an 8.9% and 5.2% lifespan increment in P4-P30 rapamycin-treated males, and 8.4% and 4.4% increment in P4-P30 treated females compared to control and P30-P60 rapamycin-treated mice, respectively (**Fig. 1C-E**). Surprisingly, P30-P60 mice did not show any significant lifespan alteration compared to control mice. This indicates that modulation of mTOR activity could influence lifespan in a specific time window, suggesting that long-term effects on lifespan can be determined early in life. P4-P30 rapamycin treatment had a profound effect on mice anatomy leading to a severe reduction in body/organ size combined with weight decrease, when compared to control mice (**Fig. 2A and fig. S1A**). Nevertheless, after P30 (the end of the treatment) the treated mice undergo a rapid body weight gain, even if they did not reach the control levels (**Fig. 2B and fig. S1B**). A significant but milder decrease in body weight was also detected during the P30-P60 rapamycin treatment (**fig. S1C**) and it was maintained after the treatment (**fig. S1D**).

**Figure 1.**
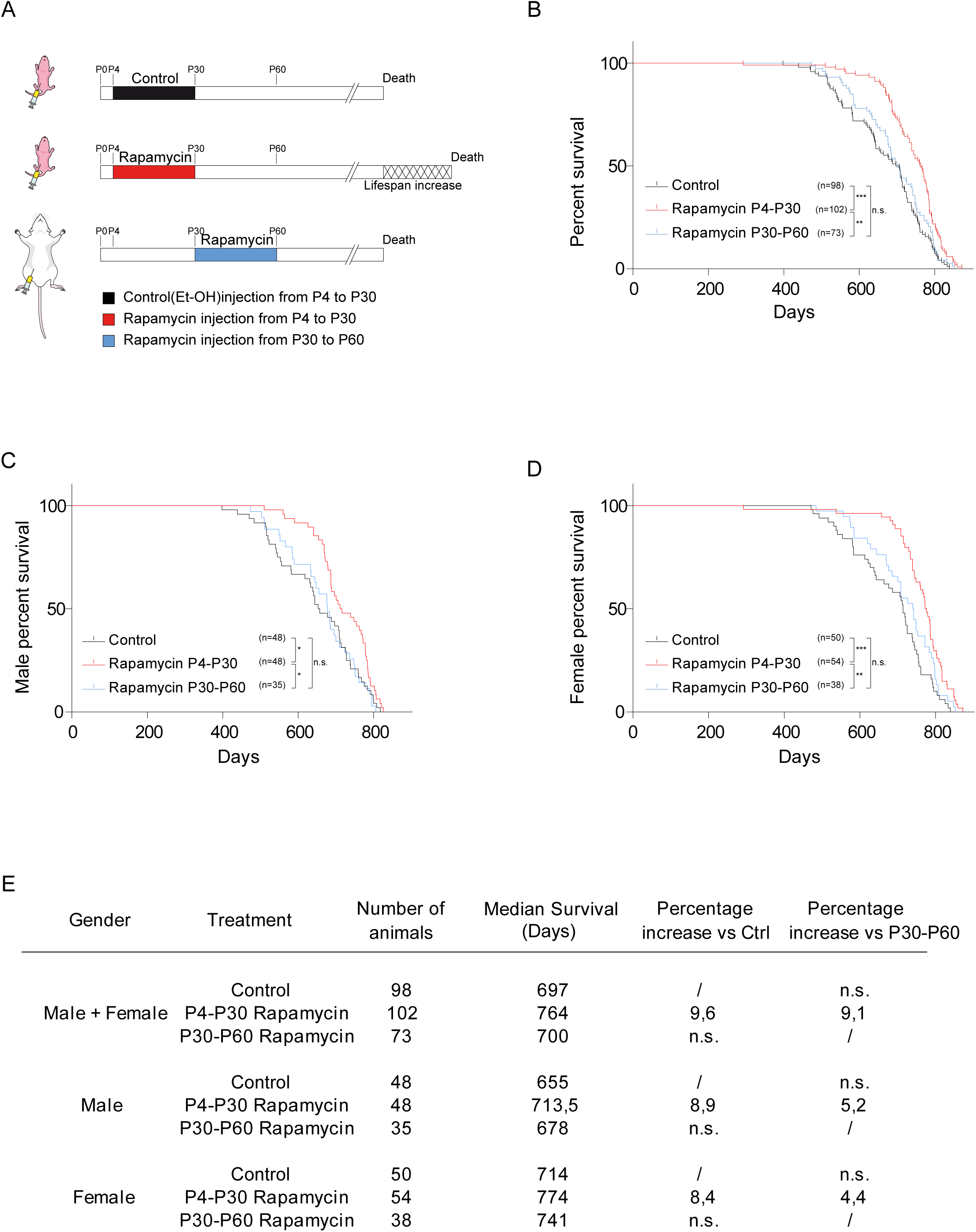
P4-P30 rapamycin treatment increases mice lifespan. **(A)** Schematic illustration of the experimental procedure and results. Control mice were intraperitoneally injected daily with Et-OH from postnatal day 4 to postnatal day 30 (P4-P30). Treated mice were intraperitoneally injected daily with rapamycin during two distinct temporal windows, P4-P30 or P30-P60. P4-P30 rapamycin treated mice show a lifespan increment compared to control and P30-P60 treated mice. **(B)** Survival curves of control mice, P4-P30 rapamycin treated mice and P30-P60 rapamycin treated mice including data from both genders (males + females). **(C, D)** Survival curves of male (**C**) and female (**D**) control mice, P4-P30 rapamycin treated mice and P30-P60 rapamycin treated mice. **(E)** Log-rank test on survival analysis (summary table).

**Figure 2.**
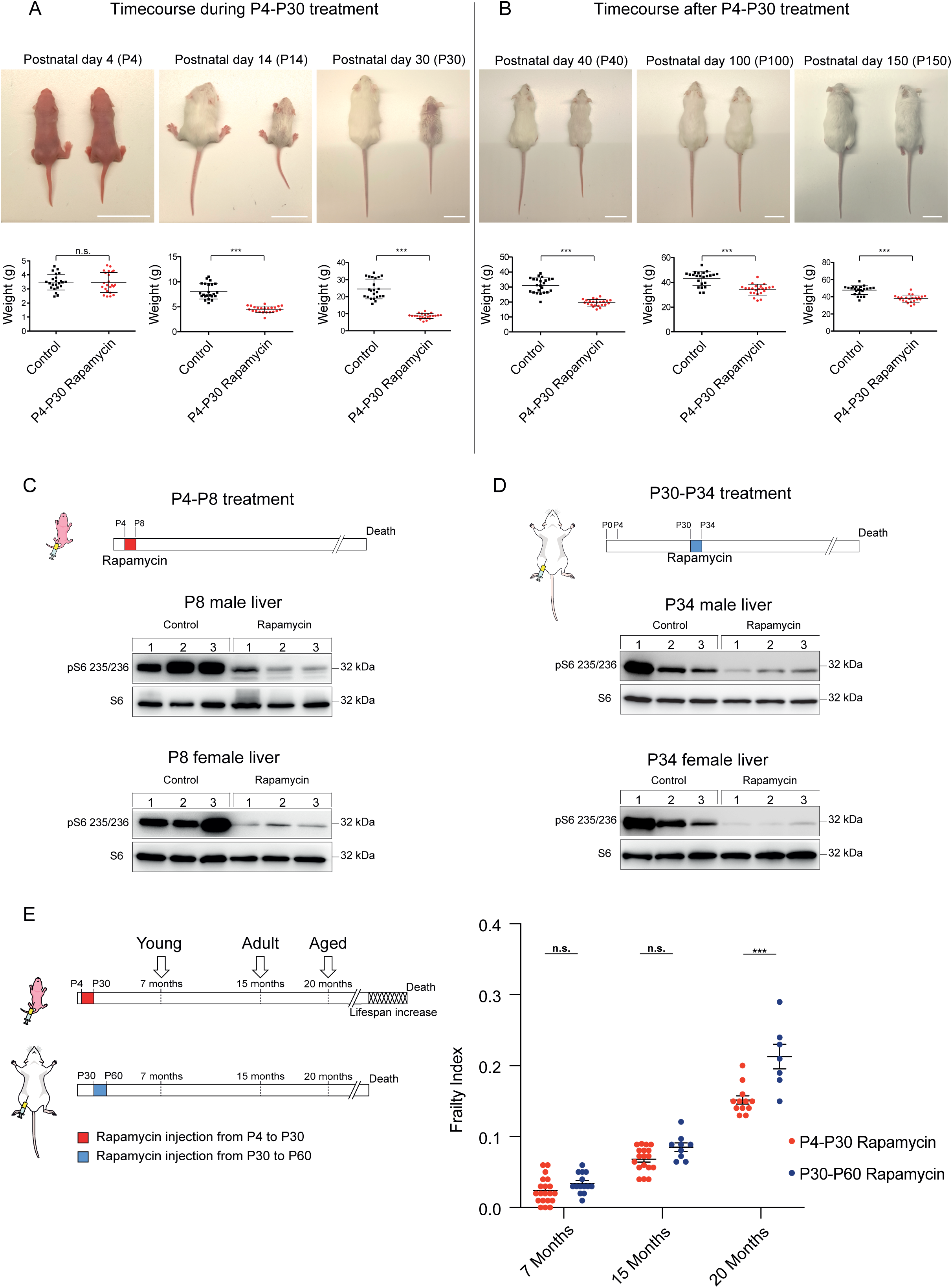
Physical and physiological status of P4-P30 and P30-P60 rapamycin treated mice. **(A)** Upper panels: Representative pictures showing rapamycin effects on mice body size during P4-P30 treatment. Images have been taken at P4, P14 and P30 (end of the treatment). Scale bar: 3 cm. Lower panels: Scatter dot plot indicating body weight of control and rapamycin treated mice during P4-P30 treatment. Data are indicated as mean + SD. **(B)** Upper panels: Representative pictures showing rapamycin effects on mice body size after P4-P30 treatment. Images have been taken at P40, P100 and P150. Scale bar: 3 cm. Lower panels: Scatter dot plot indicating body weight of control and P4-P30 rapamycin treated mice after the treatment. Data are indicated as mean + SD. **(C, D)** Upper part, schematic illustration of the experimental procedure. Western blot analysis of S6 Ribosomal Protein and phospho-S6 Ribosomal Protein (Ser235/236) from whole-liver protein extracts of female and male at P8 (**C**) and P34 (**D**) treated mice. Mice were sampled after 4 days of Et-OH or rapamycin treatment. **(E)** Schematic illustration of the different timepoints analyzed (left side) and frailty index box (right side) of P4-P30 and P30-P60 rapamycin treated mice. Red scatter dots indicate the frailty index of P4-P30 rapamycin mice at 7 months (n=20), 15 months (n=18) and 20 months (n=12). Blue scatter dots indicate the frailty index of P30-P60 rapamycin mice at 7 months (n=14), 15 months (n=9) and 20 months (n=7). Data are indicated as mean + SEM. *p-value<0.05, **p-value<0.005, ***p-value<0.0005, n.s.-not significant.

We confirmed the effectiveness of rapamycin treatment by evaluating the phosphorylation status of the ribosomal protein subunit S6 (pS6), a target substrate of S6 kinase 1 in the mTOR signaling pathway (13), in P4-P8 and P30-P34 mice livers (**Fig. 2C-D**). While mTOR inhibition is of a similar extent in the two time windows, it results in different physical and physiological long-term effects. P4-P30 rapamycin-treated mice remain smaller compared to their P30-P60 counterpart throughout life, as shown by the analysis of total length at 15 and 20 months (**fig. S1E**). Aging can be considered as a biological process determined by the accumulation of deficits that over time culminates in death. The analysis of different non-invasive parameters allows assessing a Frailty Index (FI) that can be used as a strong predictor of mortality and morbidity (18–20). P4-P30 and P30-P60 rapamycin-treated mice were monitored during all life and FI was calculated at three different time points: young, 7-months-old (210-days-old), adults,15-months-old (450-days-old) and aged, 20-months-old (611-days-old) (**Fig. 2E left side**). Although the two cohorts of mice showed no difference in the forelimb grip strength (**fig. S1F**), we observed a significant difference in the FI at 20 months between the P4-P30 and the P30-P60 treated mice. In fact, P30-P60 mice showed a worst body condition score, frequent gait disorders together with the presence of a tumor in one out of seven analyzed animals that resulted in higher FI compared to the P4-P30 rapamycin-treated mice (**Fig. 2E right side and Table S1**). Therefore, the lifespan extension of P4-P30 rapamycin-treated mice is associated with amelioration in several aging traits, resulting in a better health span compared to the P30-P60 time window. To deeper investigate the differences between the two treatments, the physical and physiological assessment has been complemented with a transcriptomic analysis of the mice liver.

To identify genes modulated by rapamycin, several groups have analyzed the hepatic gene signature in old mice subjected to continuous rapamycin treatment (21). Indeed, the liver controls several processes (i.e. hepatic glucose, insulin signaling and lipid homeostasis) potentially implicated in lifespan regulation in mammals (22, 23). Here, to identify genes and pathways involved in the lifespan extension modulated by early-life rapamycin treatment only, we analyzed by RNA-seq the gene expression profiles of P4-P30 and P30-P60 mice livers (sampled on the last day of treatment). Since it has been reported that rapamycin-mediated lifespan extension is subjected to sexual dimorphism (13, 14), we performed differential expression analysis separately for males and females (compared against the respective control samples, see methods). The differential analysis highlighted that females have fewer differentially expressed genes (DEGs) compared to the males (**Fig. 3A and Table S2**).

**Figure 3.**
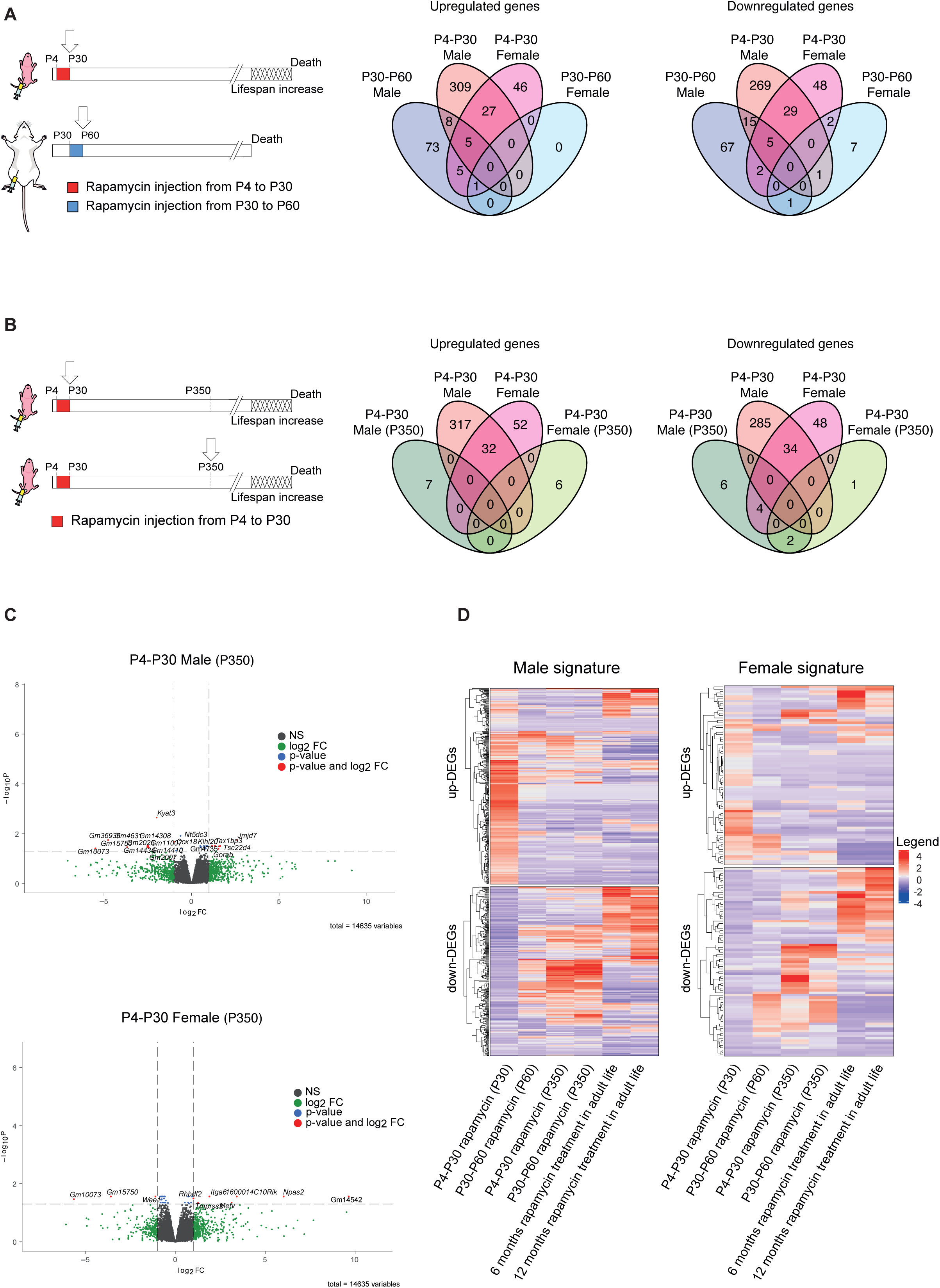
RNA-seq analysis on P4-P30 rapamycin treated mice resulted in a unique transcriptional signature. **(A)** Schematic illustration of the experimental procedures (left). Landscape of up- and down-regulated genes across P4-P30 and P30-P60 treatments in male and female mice. Venn diagrams are used to highlight private and shared differentially expressed genes. **(B)** Schematic illustration of the experimental procedures (left). Landscape of up- and down- regulated genes across P4-P30 treatment processed at the last day of treatment and at P350 in male and female mice. Venn diagrams are used to highlight private and shared differentially expressed genes. **(C)** Volcano plots showing the transcriptional changes in P4-P30 rapamycin treated male (upper panel) and female (lower panel) treated mice processed at P350. The log2FC is represented on the x-axis. The y-axis shows the -log10 of the corrected p-value. A p-value of 0.05 and log2FC of 1 and -1 are indicated by gray lines. Top 10 upregulated and top 10 downregulated genes (when available) are labeled with gene symbols. **(D)** Log2FC of genes that are differentially expressed only in male (left side) and female (right side) in response to P4-P30 treatment at the last day of treatment are compared, through heatmaps, with corresponding log2FC profiles in: P30-P60 on the last day of treatment; P4-P30 and P30-P60 treatment analyzed at P350; published data on chronic rapamycin treatment in adult life (6 and 12 months) (ref.21).

To understand the long-term effects of early rapamycin treatments we analyzed the transcriptome landscape of P4-P30 and P30-60 mice at the middle life stage (P350). Interestingly, at P350 we observed an absence of significant transcriptome differences, with only a few mildly deregulated genes (**Fig. 3B-C and Table S3**). These results confirm the concept that the inhibition of mTOR leads to different effects depending on the time of the inhibition (24). In addition, we investigated whether the P4-P30 rapamycin associated signature correlates with the already published signature derived from chronic treatments. To do so, we compared P4-P30 RNA-seq data with the dataset obtained from two different chronic treatments starting at 4 months of life that differ for the administered dose of rapamycin (42 ppm and 14 ppm, respectively) and treatment duration (6 and 12 months, respectively) (21). The analysis showed that the early transient rapamycin treatment possesses a unique gene expression signature (**Fig. 3D**) leading us to focus on the gene expression changes on the last day of treatment.

To identify pathways associated with the altered gene signatures, we performed Gene Set Enrichment Analysis (GSEA) (**Fig. 4A and fig. S2A**). The analysis revealed that the two time windows of treatment lead to different and at times divergent gene set enrichments (**Fig. 4A**). For example, P4-P30 and P30-P60 treatments show opposite effects on chromatin binding and oxidoreductase activity (**Fig. 4A and fig. S2A and Table S4**). Overall, the analysis highlighted the broad upregulation of the sulfotransferase gene family by rapamycin treatment. In fact, we observed an enrichment of several molecular functions related to sulfotransferase activity (steroid-, bile salt-, and alcohol-sulfotransferase activity) in P4-P30 treated males that is not present in the P30-P60 time window (**Fig. 4A**). Interestingly, among the common deregulated genes in P4-P30 males and females, the sulfotransferases *Sult2a3* and *Sult2a6* emerged as two of the upregulated genes and *Sult5a1* as downregulated gene (**Fig 4B and Table S2**). Moreover, several sulfotransferases (SULTs), such as *Sult1d1, Sult1e1, Sult2a1, Sult2a2, Sult2a3, Sult2a4, Sult2a5, Sult2a6, Sult2a8* and *Sult5a1* are upregulated only in the P4-P30 time window (**Fig. 4C**). Sulfotransferases are a super gene family of enzymes involved in sulfonate conjugation processes, catalyzing the transfer of sulfonate (SO_3_^-^) to a hydroxyl or amino group. Hepatic regulation and activity of SULTs could vary based on age and sex (25) and their expression is controlled by numerous members of the nuclear receptor (NR) superfamily, that act as sensors of xenobiotics as well as endogenous molecules, such as fatty acids, bile acids, and oxysterols (26). To determine whether the functional enrichment in sulfotransferase activity can be linked to metabolic remodeling, we performed hepatic metabolomics analysis on P4-P30 and P30-P60 rapamycin-treated mice. A total of 138 metabolites were detected in the P4-P30 and P30-P60 time windows. We identified 70 metabolites that were significantly different across the two treatments (**Table S5**). The P4-P30 rapamycin treatment revealed a unique metabolic signature (**Fig. 5A-B**) and the deregulation of several primary and conjugated bile acids (**Table S5**). Of notice, bile acids biosynthesis was identified as one of the most enriched and significant pathways by the Metabolic Set Enrichment Analysis (MSEA) (**Fig. 5C**). Most of the detected primary and conjugated bile acids are downregulated in P4-P30 rapamycin-treated mice compared to the P30-P60 time window, suggesting the upregulation of the detoxifying pathway (**Fig. 5D**). This observation is consistent with previous findings where higher detoxification capacity was associated with reduced intrahepatic bile acid levels (27). The same analysis on P4-P30 and P30-P60 control mice did not show the same results, indicating only a few deregulated metabolites and a stronger overlap between the two time windows (**Fig. S3A-C and Table S5**). Interestingly, the metabolic differences revealed on the last day of treatment are not maintained later in life. Indeed, the same analysis on P4-P30 and P30-P60 at P350 did not reveal differences across the two treatments, showing overlap among the two time windows (**Fig. 5E-F, fig. S3D-E and Table S5**). Together, these results suggest that metabolic data correlate with the transcriptome analysis, leading to focus on the changes at the last day of treatment, that reflect the direct effect of lifespan-extending intervention, and indicating sulfotransferase activity as an important process affecting the early and transient rapamycin treatment at P4-P30.

**Figure 4.**
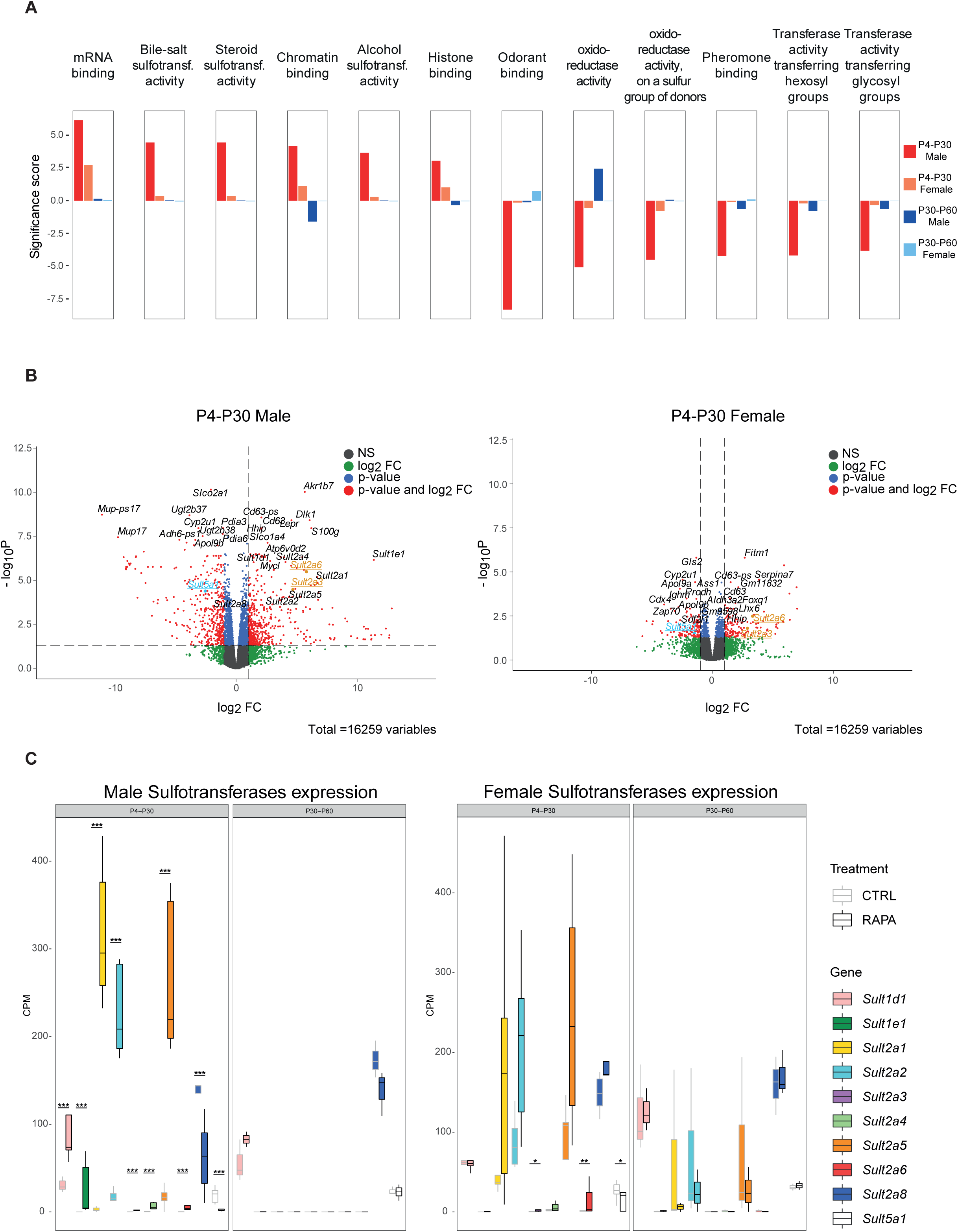
Functional enrichment analysis indicates sulfotransferases activity as a pathway involved in lifespan regulation. **(A)** Gene set enrichment analysis (GSEA) results of P4-P30 and P30-P60 at the last day of treatment in male and female mice. Significance score, calculated as -log10(q-value) corrected by the sign of regulation, is plotted on the y axis. Plots are representative of the top 12 GO Molecular Function (MF) terms with higher/lower significance scores for the male P4-P30 rapamycin treated mice (top 6 with higher significance scores and top 6 with lower significance scores). The whole list of enriched GO terms is available in Table S4. **(B)** Volcano plots showing the transcriptional changes in P4-P30 rapamycin treated male (left side) and female (right side) treated mice. Each circle represents a gene. Underlined and highlighted terms are common *SULTs* genes shared by males and females (orange for upregulated genes, blue for downregulated genes). The log2FC is represented on the x-axis. The y-axis shows the -log10 of the corrected p-value. A p-value of 0.05 and log2FC of 1 and -1 are indicated by gray lines. Top 10 upregulated and top 10 downregulated genes are labeled with gene symbols. **(C)** Expression profile of sulfotransferases deregulated only in the P4-P30 time window in male (left side) and female (right side) treated mice. Values for treated (dark borders) and control (light borders) samples across the different conditions are shown as median CPM with bars representing standard deviations across the biological replicates. *p-value<0.05, **p-value<0.005, ***p-value<0.0005.

**Figure 5.**
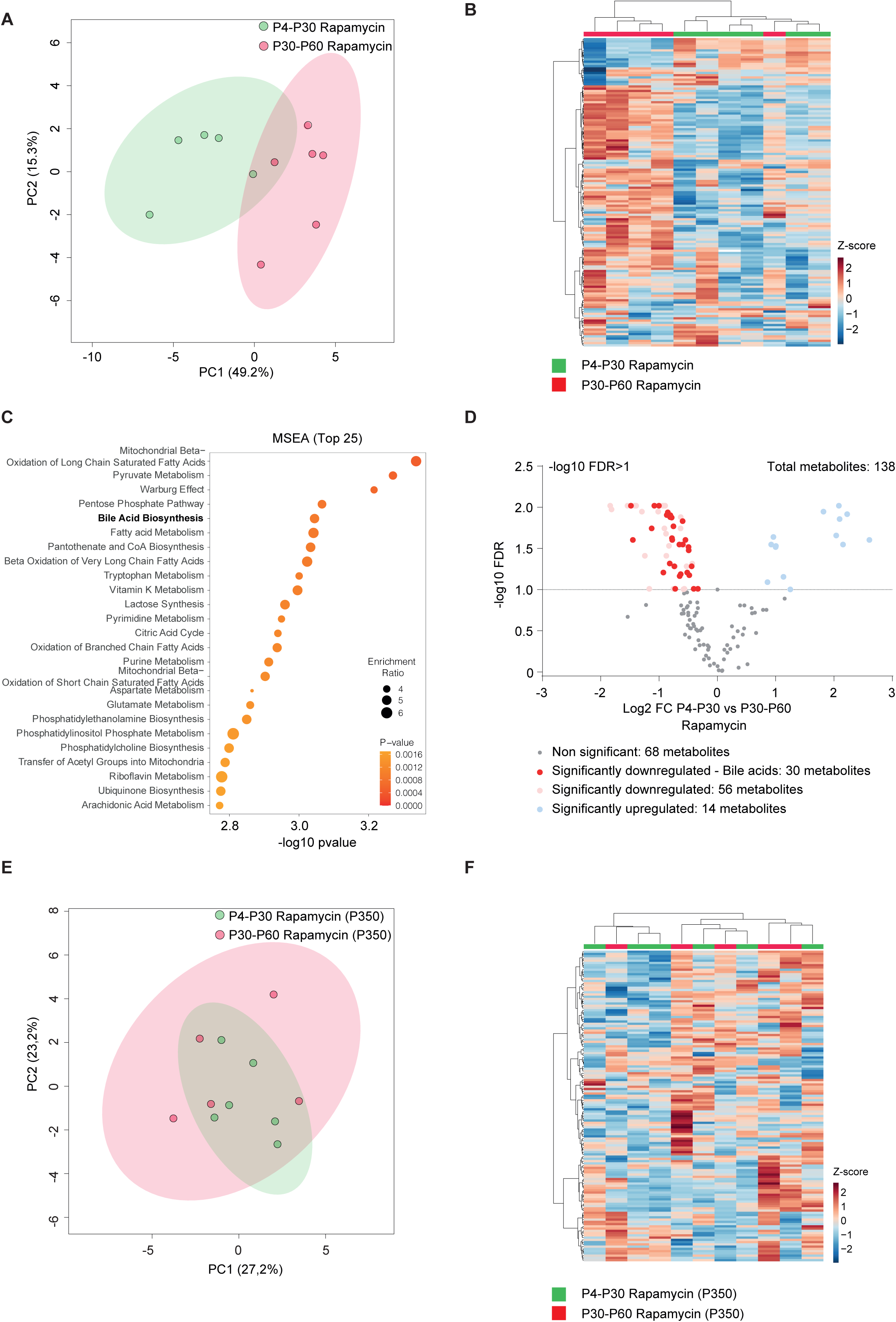
Early transient treatment with rapamycin increases bile acid liver detoxification in mice. **(A, B)** Principal Component Analysis (PCA) (**A**) and heatmap (**B**) of liver metabolomic profile from P4-P30 (green samples) and P30-P60 (red samples) mice treated with rapamycin. **(C)** Top 25 metabolic pathways enriched in P4-P30 compared to P30-P60 mice treated with rapamycin. Metabolic Set Enrichment Analysis (MSEA) was performed taking advantage of MetaboAnalyst 5.0 webtool interrogating KEGG database. The x-axis shows the -log10 of *p*value. **(D)** Volcano plots showing the metabolomic changes in P4-P30 compared to P30-P60 mice treated with rapamycin. Each circle represents one metabolite. The log2 fold change is represented on the x-axis. The y-axis shows the -log10 of the False Discovery Rate (FDR). A FDR of 0.1 is indicated by gray line. Grey, pink and light blue dots represent unchanged, significantly downregulated and significantly upregulated metabolites, respectively. Red dots represent significantly downregulated bile acids in P4-P30 compared to P30-P60 mice treated with rapamycin. **(E, F)** Principal Component Analysis (PCA) (**E**) and heatmap (**F**) of liver metabolomic profile from P4-P30 (green samples) and P30-P60 (red samples) mice transiently treated with rapamycin and analyzed at P350.

To clarify the role of rapamycin in the modulation of sulfotransferases we decided to investigate the gene expression profiles of more acute treatments, such as P4-P8 and P30-P34. The inhibition of the mTOR pathway has a stronger transcriptome effect early in life (P4-P8), while only a few genes are deregulated in expression when the treatment occurs later (P30-P34) (**fig. S2B**). Interestingly, the nuclear hormone receptor *Nr1i3* (*CAR*) is upregulated in both male and female P4-P8 (**fig. S2C and Table S2**). CAR regulates the expression of several sulfotransferases (26), and its upregulation upon rapamycin treatment might explain the enrichment of this category of genes in the P4-P30 treated mice. This data strongly supports the idea that rapamycin has different effects based on the age of administration and suggest that the sulfotransferase pathway may be involved in lifespan regulation.

The discovery that lifespan can be regulated in the early stage of life in mice prompted us to investigate whether this process was evolutionarily conserved. We recreated similar experimental conditions as in mice, by administering rapamycin during larval development of *Drosophila melanogaster*. *Drosophila* life cycle can be divided into 4 developmental stages: embryo, larva (three instar stages), pupa and adult that correspond to four distinct periods of life: embryonic development, a juvenile growth phase, sexual maturation, and reproductive adulthood, respectively (28). *Drosophila* growth occurs mainly during the juvenile larval stages, and the transition between the second (L2) and third larval instar (L3) represent an important time window during which the animal reaches the “critical weight” to continue the development (29). For this reason, rapamycin treatment was carried out in the isogenic *white iso31* (herein *w^iso31^*) *Drosophila* strain during the third instar larvae stage. *w^iso31^* third instar larvae were treated with 1 µM, 50 µM or 200 µM rapamycin, starting from 72 hours after egg-laying till pupal stage (**Fig. 6A upper timeline and fig. S4A upper timeline**). Although flies exposed to 1 µM and 50 µM do not show a significant increase in lifespan for both sexes (**Fig. S4 B-G and Table S6**), treatment with 200 µM rapamycin led to an increase in lifespan compared to the control (**Fig. 6B**). As observed in mice, the treatment led to a reduction in body size that is maintained during all the developmental stages (**fig. S5A**). Early transient rapamycin treatment on *Drosophila* larvae determined a significant increase in median lifespan when both genders were analyzed together (**Table S6**), while, when the genders were analyzed separately only male flies showed a significant lifespan extension compared to control flies (**Fig. 6B-D**). To verify whether the modulation of mTOR activity influences *Drosophila melanogaster* lifespan only in a specific time window as for the mammalian counterpart (**Fig. 1A, B**), we tested the effect of rapamycin treatment at later timepoint. When 10-days-old *w^iso31^* flies were treated with 200 µM rapamycin for 3 days (**Fig. 6A lower timeline**) no effect was observed in terms of body size and lifespan compared to control (**Fig. 6E-G and fig. S5B**). The effectiveness of rapamycin treatment in the two time windows was evaluated by profiling the phosphorylation status of S6K by western blot analysis, using a phospho-Thr398-dependent S6K antibody (14). Flies exposed to 200 µM rapamycin at the last day of treatment (wondering larvae and 13-days-old, respectively) showed a comparable reduction in phospho-T398-S6K, suggesting that TOR signaling is similarly down-regulated in the two time windows (**fig. S5C**). These results strengthen the idea that lifespan can be determined by transient early-life events and that similar treatments at later time points have no effect on lifespan. Although the mechanism is evolutionarily conserved, *Drosophila* and mouse development is significantly different and difficult to compare. To investigate if TOR inhibition could influence *Drosophila* lifespan during other time windows, we treated *w^iso31^* flies with 200 µM rapamycin during early stages of adult life, such as the first 10 days after eclosion (adult emergence from the pupal cases) (**Fig. 6A**). Interestingly, rapamycin-treated flies displayed an increase in lifespan compared to the control (**Fig. 6H**) leading to an increment in the median lifespan for both sexes (**Fig. 6I-J and Table S6**). Moreover, to verify that the effect was due to the specific time window of administration and not to the duration of the treatment, we also treated 10-days-old *w^iso31^*flies with 200 µM rapamycin for 10 days (**Fig. 6A**) and we observed no effect on lifespan (**Fig. 6K-M and Table S6**). In conclusion, our results indicate that inhibition of TOR in specific time windows early in life is an evolutionary conserved mechanism that leads to lifespan extension.

**Figure 6.**
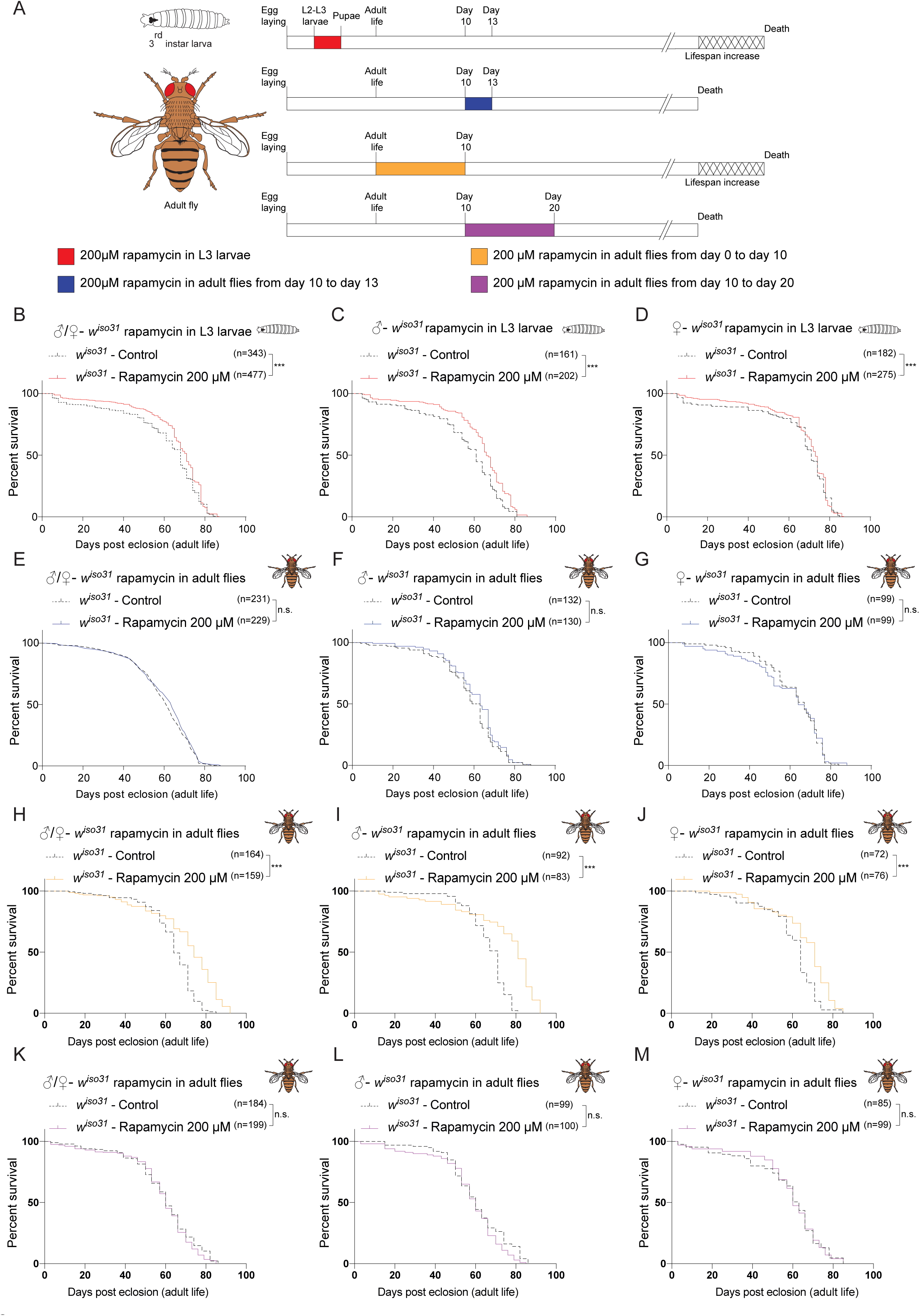
Early transient rapamycin treatment on *w^iso31^ Drosophila melanogaster* induces a time-dependent effect on lifespan. **(A)** Schematic illustration of the experimental procedure and results. Flies were transiently treated during larval stages with rapamycin 200 µM starting from 72 hours after egg-laying to puparium formation (red) or during adulthood, from day 0 to day 10 (orange), from day 10 to day 13 (blue) or from day 10 to day 20 (purple). Rapamycin administration during development and during the first 10 days of life, but not at later timepoints, leads to lifespan increment. **(B)** Survival curves of *w^iso31^* flies transiently treated from 72 hours after egg-laying till puparium formation (males + females) with Et-OH (control) or rapamycin 200 µM. **(C, D)** Survival curves of male (**C**) and female (**D**) *w^iso31^*flies transiently treated from 72 hours after egg-laying till puparium formation with Et-OH (control) or rapamycin 200 µM. **(E)** Survival curves of *w^iso31^* flies transiently treated in adult stage, from day 10 to 13 (males + females), with Et-OH (control) or rapamycin 200 µM. **(F, G)** Survival curves of male (**F**) and female (**G**) *w^iso31^*flies transiently treated in adult stage, from day 10 to 13, with Et-OH (control) or rapamycin 200 µM. **(H)** Survival curves of *w^iso31^* flies transiently treated from day 0 to day 10 (males + females) with Et-OH (control) or rapamycin 200 µM. **(I, J)** Survival curves of male (**I**) and female (**J**) *w^iso31^*flies transiently treated from day 0 to day 10 with Et-OH (control) or rapamycin 200 µM. **(K)** Survival curves of *w^iso31^* flies transiently treated from day 10 to 20 (males + females), with Et-OH (control) or rapamycin 200 µM. **(L, M)** Survival curves of male (**L**) and female (**M**) *w^iso31^*flies transiently treated from day 10 to 20, with Et-OH (control) or rapamycin 200 µM. *p-value<0.05, **p-value<0.005, ***p-value<0.0005, n.s.- not significant.

As previously described, the RNA-seq experiment performed on mice liver (**Fig. 4A-B**), suggested that sulfotransferases could be involved in lifespan extension induced by the rapamycin treatment. *Drosophila* harbors four *SULTs* orthologues, namely *dST1*, *dST2*, *dST3* and *dST4,* thought to be derived from gene duplication processes occurred in a common ancestral gene. Among them, *dST1* and *dST3* show a high degree of homology (30, 31) and as shown in **fig. S6A-B,** we observed increased *dST1* and *dST3* mRNA levels upon 12 hours of rapamycin treatment during larval development, but not in adult (10-days-old) treated flies, thus supporting the idea that rapamycin has different effects based on age of administration, as observed in mammals. Of notice, dST1 (CG5428) shares 46% and 47% of similarity in the amino acidic sequence with the mouse Sult2a3 and Sult2a6 respectively counterparts that are upregulated in both male and female P4-P30 rapamycin treated mice (**Fig. 4B**). To investigate the role of *dST1* in lifespan extension, we took advantage of the GAL4-UAS system (32) to induce *dST1* constitutive overexpression during the entire life of flies, using a *TubGal4* promoter (**Fig. 7A upper timeline).** Lifespan was compared to a transgenic strain carrying the same genetic weight (*UAS-GFP;tubGal4/+*). Constitutive *dST1* upregulation in transgenic flies determines a decrease in lifespan compared to control flies of both sexes (**Fig. 7B-D**). Since prolonged upregulation seems to be detrimental, we aimed to recreate the same experimental conditions of rapamycin treatment by transiently overexpressing *dST1* during larval development only (**Fig. 7A lower timeline**). Transient *dST1* upregulation was achieved using a *tubGal80^TS^;TubGal4* strain, and the lifespan was compared to a transgenic strain carrying the same genetic weight (*tubGal80^TS^/UAS-GFP;tubGal4/+*). *dST1* overexpressing flies displayed an extension of lifespan compared to control (**Fig. 7E and Table S6**) and the effect was present in both males and females (**Fig. 7F-G and Table S6)**. These data support the previous rapamycin experiments on mice and *Drosophila* (**Fig. 1A and Fig. 6A**) and confirmed the presence of precise time windows during which lifespan can be affected. To characterize the effect of transient *dST1* overexpression on the fitness of the flies, we tested their endurance by analyzing the motor function during aging with a negative geotaxis assay. Indeed, locomotor behavior is considered as a marker of organismal health that declines with age (8, 14). Early and transient *dST1* up-regulation in larvae improved the climbing performance over time compared to controls (**Fig. 7H**) and the amelioration was present in both sexes (**Fig. 7 I-J**). Overall, our novel findings demonstrate that lifespan can be determined during early life, unveiling the role of *dST1* as a new lifespan modulator.

**Figure 7.**
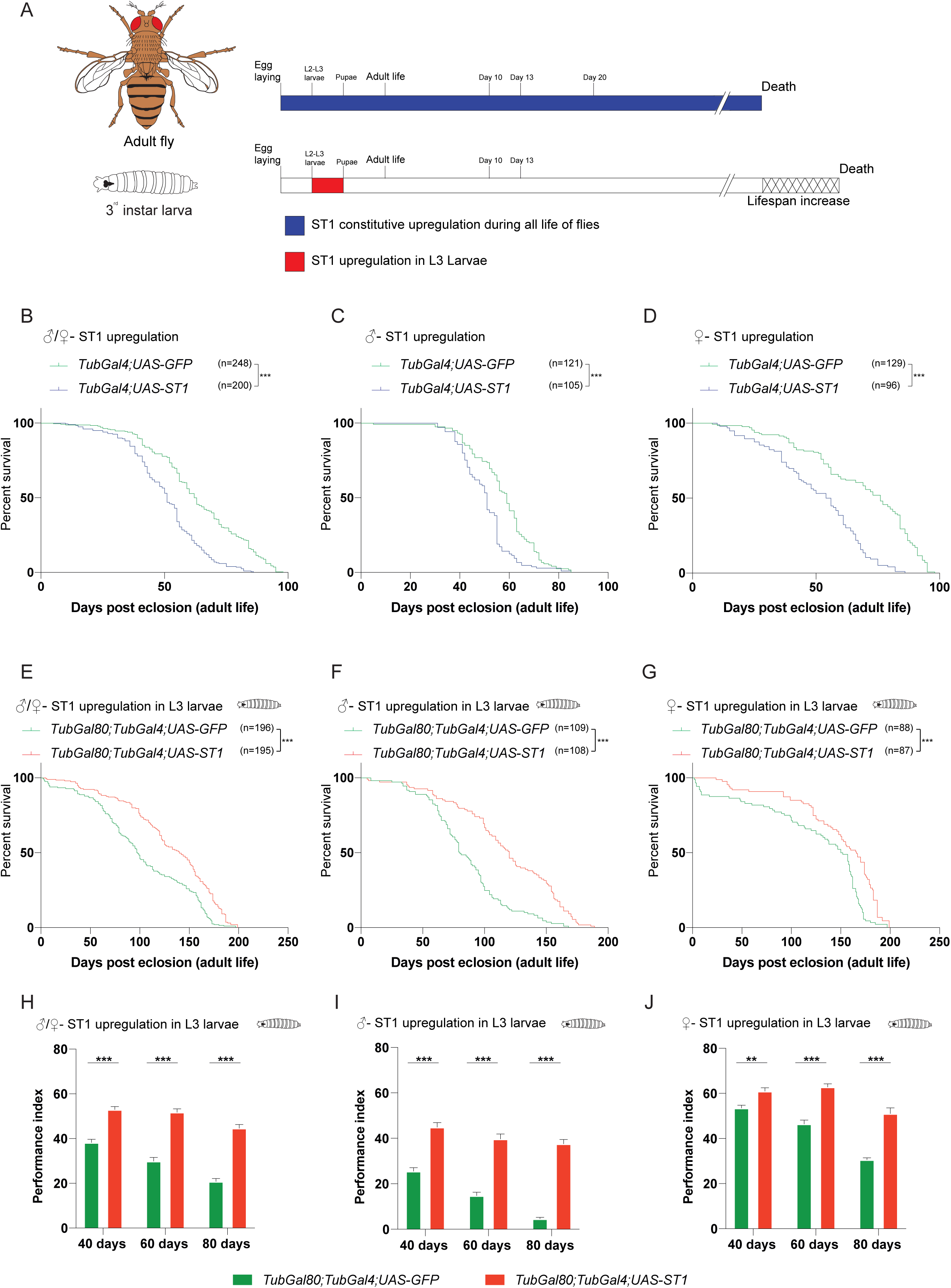
Transient dST1 overexpression during larval stage, determines an increase in *Drosophila* m*elanogaster* lifespan. **(A)** Schematic illustration of the experimental procedure and results. Constitutive dST1 overexpression during the entire life (blue) of flies do not increase lifespan. Transient dST1 overexpression during larval stage (red), leads to lifespan increment. **(B)** Survival curves of flies harboring constitutive dST1 upregulation (*TubGal4/UAS-dST1)* compared to control strain (*UAS-GFP;TubGal4/+)* males + females. **(C, D)** Survival curves of flies harboring constitutive dST1 upregulation (*TubGal4/UAS-dST1)* in male (**C**) and female (**D**) compared to control strain (*UAS-GFP;TubGal4/+)*. **(E)** Survival curves of flies upon upregulation of dST1 during larval stages (*tubGal80^TS^/+;TubGal4/UAS-dST1)* compared to the control strain *tubGal80^TS^/UAS-GFP;TubGal4/+* (males + females). **(F, G)** Survival curves of male (**F**) and female (**G**) flies upon upregulation of dST1 during larval stages in (*tubGal80^TS^/+;TubGal4/UAS-dST1)* compared to control strain *tubGal80^TS^/UAS-GFP;TubGal4/+*. (H) Climbing performance index of *tubGal80^TS^/UAS-GFP;TubGal4/+* and *tubGal80^TS^/+;TubGal4/UAS-dST1* at different timepoints: 40 days (n=176, 196 respectively), 60 days (n=159, 178 respectively) and 80 days (n=119, 158 respectively). Upregulation of dST1has been induced only during larval stages. (I) Climbing performance index of male *tubGal80^TS^/UAS GFP;TubGal4/+* and *tubGal80^TS^/+;TubGal4/UAS-dST1* at different timepoints: 40 days (n=93, 93 respectively), 60 days (n=79, 81 respectively) and 80 days (n=41, 70 respectively). Upregulation of dST1has been induced only during larval stages. (J) Climbing performance index of male female *tubGal80^TS^/UAS-GFP;TubGal4/+* and *tubGal80^TS^/+;TubGal4/UAS-dST1* at different timepoints: 40 days (n=83, 103 respectively), 60 days (n=80, 97 respectively) and 80 days (n=78, 88 respectively). Upregulation of dST1has been induced only during larval stages. *p-value<0.05, **p-value<0.005, ***p-value<0.0005, n.s.-not significant).

## Discussion

Environmental and genetic components influence lifespan by regulating specific signaling pathways, metabolism, and transcription factors. So far, the great majority of studies were based on pharmacological treatments administered late in life for a prolonged time or repeated treatments. On the other hand, only a few groups studied how perturbation in the early life of mice and flies (i.e., caloric restriction/modulation or antioxidant treatments) affects lifespan (3,5,8). Our results indicate a critical early-life timeframe during which the modulation of age-related pathways determines a long-term effect on lifespan. By exploiting a transient rapamycin treatment, we identified a crucial lifespan-extending time window both in mouse (P4-P30) and in *Drosophila melanogaster* (larval stage and early adult).

Interestingly, the transient inhibition of the mTOR pathway in later periods of life does not significantly improve the lifespan. Our results suggest the existence of a ‘memory’ mechanism that increases lifespan, and that can be modulated in early life only. A similar hypothesis has been postulated in mice exposed to caloric restriction in adult life (33). These mice showed signs of a ‘nutritional memory’ and metabolic remodeling of white adipose tissue, but the molecular mechanisms beyond these effects are unknown (33).

To identify genes that affect lifespan, we analyzed gene expression profiles of livers from mice transiently treated with rapamycin early in life (P4-P30). We chose to profile the liver, because this tissue regulates glucose, insulin signaling and lipid homeostasis and could potentially regulate mammalian lifespan (22, 23). Furthermore, hepatic gene signatures of different lifespan-extending treatments have been already used to identify aging-related candidate genes (21). Indeed, P4-P30 rapamycin-treated mice processed on the last day of treatment show a unique and distinct hepatic gene signature, different from the other transient treatments. Of notice, this signature is not maintained during later time points. Indeed, gene expression changes cannot be observed in middle life (P350). These results led us to speculate that the P4-P30 modulation of the mTOR pathway determines a chain of events set in motion during the early-life time window, revealing new age-related genes with novel functions in regulation of lifespan. Importantly, GSEA analysis on P4-P30 treated mice (males) resulted in the enrichment of several molecular functions related to sulfotransferase activity (steroid-, bile salt-, and alcohol-sulfotransferase activity) while *Sult2a3* and *Sult2a6* were found upregulated in both sexes. *SULTs* are generally involved in the xeno- and endobiotics metabolism that is divided into three phases: (I) modification, (II) conjugation and (III) excretion. Those enzymes catalyzed the transfer of sulfonate (SO_3_^-^) to a hydroxyl or amino-group, favoring the elimination from the body (34). Sulfonation occurs on numerous xeno- and endobiotics such as drugs, steroid hormones, bile acids, peptides and lipids and it has been generally considered as a detoxification pathway generating an end-product that is more amenable to eliminate (34). Xenobiotic metabolism has been already associated with lifespan extension in *Drosophila melanogaster*. Indeed, early low doses of oxidants determine a long-term mechanism that leads to lifespan extension (7). However, the constitutive upregulation of xenobiotic resistance mediators correlates with health span amelioration, but not with lifespan extension (35). For this reason, we decided to test the early transient lifespan-extension role of sulfotransferase taking advantage of *Drosophila melanogaster*, that has a considerably shorter lifespan than mice and it is characterized by the presence of genetic tools that allow temporal modulation of gene expression. Indeed, transient *dST1* overexpression (*Sult2a3* and *Sult2a6 Drosophila* homologs) during larval development, determines a significant increase of lifespan revealing a new putative function of *dST1* as an early regulator of the aging process. Our findings indicate a link between early-life events and long-term effects on lifespan indicating the existence of a critical time window that can permanently affect how long an individual can live. We found that the modulation of gene expression/pathways in this specific time window can determine lifespan extension both in mice and *Drosophila melanogaster*. Further studies are needed to assess the role of sulfotransferases and their regulators in the aging process and to unveil new drugs that might increase lifespan through an early and transient administration. However, our data represent a new starting point for the study of lifespan, paving the path for future work on humans.

## Acknowledgments

We thank all members of the Tiberi laboratory for technical expertise and feedback. We thank Ilaria Morassut, Dr. Alessandro Alaimo and Dr. Annalisa Rossi for helpful discussion and advices. We thank Sergio Robbiati (MOF facility) and Veronica De Sanctis, Roberto Bertorelli, and Paola Fassan (NGS facility). We thank Dr. Alex Gould for providing us with the *white iso31 Drosophila melanogaster* isogenic strain. We thank the Zurich ORFeome Project for the *Drosophila* stocks.

## Funding

G.A. was supported by a FIRC-AIRC fellowship for Italy. F.A. was supported by Fondazione Umberto Veronesi post-doctoral fellowship (il Dono di Rossana) and Marie Skłodowska Curie fellowship (grant agreement 844677). This work was funded by a grant from the Giovanni Armenise-Harvard Foundation, United States (Career Development Award 2016, to L.T.) and My First AIRC Grant, Italy (Project Code: 19921 to L.T. and 20621 to A.R), EMBO grant to L.T.

## Author contributions

G.A. and L.T. designed the study, analyzed data, and wrote the manuscript; A.S. supervised the *Drosophila melanogaster* experiments; F.A., A.S. and A.Q. helped for manuscript revision; G.A. performed all the experiments with help from C.S., M.G., and A.S.; G.A. and C.S. performed *in vivo* experiments; Bioinformatics analyses were performed by D.P and A.R.; Metabolomic data and analyses were performed by M.A. and N.M.

## Competing interests

The authors declare no competing interests.

## Materials and Methods

### Mice

CD1 mice (cat. #022) were purchased from Charles River and housed in a certified specific pathogen-free (SPF) Animal Facility in accordance with European Guidelines. Mice were provided ad libitum food access throughout their lifetime. Male and female CD1 mice were treated with rapamycin or ethanol at different time points and used for the survival analyses. Mice were monitored daily until human endpoint, caused by death or euthanasia due to the occurrence of severe age-related pathologies, identified by veterinary and biological services staff members. Survival analysis was performed using mice born within the same period (3-4 months) and all of them derived from a small cohort of male and female CD1 mice, thus decreasing the differences in the genetic background The experiments were approved by the Italian Ministry of Health as conforming to the relevant regulatory standards.

### Rapamycin treatment and survival analysis in mice

Rapamycin (Alfa Aesar, cat. #J62473) was dissolved in ethanol at 20 mg/mL and then diluted in milli-Q water. CD1 mice were daily intraperitoneally injected with rapamycin (10 mg/kg) or ethanol for two distinct time windows from P4 to P30 or from P30 to P60 and sacrificed at the end of treatments for RNA-seq analysis or monitored until human endpoint for the survival analysis. Acute rapamycin effect was evaluated intraperitoneally injecting CD1 mice with either rapamycin (10 mg/kg) or ethanol from P4 to P8 or from P30 to P34 and sacrificed at the end of treatments for Western Blot and RNA-seq analysis. Control and rapamycin treated animals were defined per cage to ensure similar mother feeding among the rapamycin treated mice. Control and rapamycin treated mice were weaned at the same age.

### Weight analysis

Rapamycin treated and control CD1 mice were weighed daily in the morning to check weight changes during and after rapamycin treatment. Weight measurements were collected from all the different time-windows: P4-P30 control (n=3), P4-P30 rapamycin (n=2), P30-P60 control (n=3), P30-P60 rapamycin (n=3). Mean weight measurements of each litter was calculated considering both males and females.

### Length analysis

Measurements have been performed on P4-P30 and P30-P60 rapamycin treated mice at 15 months and 20 months. Mean length measurements was calculated considering both males and females.

### Western Blot

Mouse section: Whole livers of mice subjected to four days of treatment (P4-P8 and P30-P34) were dissected and snap frozen from males and females CD1 mice at P8, and P34 respectively. Fresh-frozen livers were smashed with mortar and pestle and proteins were extracted from smashed tissues in lysis buffer (50mM Tris-HCl, 150mM NaCl, 20mM EDTA, 1% NP-40, 0,5% sodium deoxycholate, 0,1% SDS), supplemented with proteases inhibitors (VWR, cat. #M221-1ML), DTT (Thermo Fisher Scientific, cat. #R0861), and Serva Electrophoresis™ Phosphatase-Inhibitor-Mix II Solution (Thermo Fisher Scientific, cat. #3905501). *Drosophila* section: L3 wondering larvae (n=30) and 10-days-old flies (n=10) treated for three days with rapamycin or ethanol, were collected and homogenized using a pellet pestle (Sigma-Aldrich, cat. #Z359971-1EA) to obtain a better homogenization of the larvae and adult flies whole bodies in the same lysis buffer used for mouse whole liver extracts (see above). Samples in lysis buffer were left in ice for 30 min and then centrifuged at 18000 g for 20 min. Supernatants, containing extracted proteins, were collected in new eppendorf tubes and proteins were quantified using the Bradford method and stored at -80 °C. Proteins were resolved by SDS-PAGE and transferred onto a PVDF membrane (pore size 0,2 μm, Merck, cat. #GE10600021. The membrane was blocked in 5% BSA (Thermo Fisher Scientific, cat. #11423164)/TBS-T (0,1% Tween in TBS) for 1 h at room temperature and low agitation, and subsequently probed with primary antibodies, diluted in 5% BSA/TBS-T, overnight at 4 °C. Then, the membrane was washed with TBS-T three times for 10 min and incubated with secondary antibodies, diluted in 5% BSA/TBS-T, for 1 h at room temperature. After another cycle of three washes with TBS-T, protein levels were detected using the Clarity Western ECL Substrate (Biorad, cat. #1705062). The harsh stripping protocol was applied to detect the phosphorylation state of protein in the same PVDF membrane. To remove primary and secondary antibodies, the membrane was incubated with a stripping buffer (20 mL 10% SDS, 12,5 mL 0,5M Tris-HCl pH 6.8, 67,5 mL distilled water supplemented with 0,8 mL β-mercaptoethanol (Scharlab, cat. # ME00950250)) at 50 °C for 45 min. Then, the membrane was rinsed with milli-Q water for 1-2 min and with TBS-T for 5 min. After another step of blocking the membrane was incubated with a new primary antibody.

### RNA-extraction, library generation and sequencing

P4-P8 (n=3), P4-P30 (n=5), P30-P34 (n=3), P30-P60 (n=3) and P350 (n=3) control and rapamycin treated whole livers were dissected and snap frozen from males and females CD1 mice at P8, P30, P34, P60 and P350 respectively. Fresh-frozen livers were smashed with mortar and pestle and total RNA was isolated from smashed tissues with TRIzol Reagent (Invitrogen, cat. #15596018), according to the manufacturer’s instructions. Pellet pestles (Sigma-Aldrich, cat. #Z359971-1EA) were used to obtain a better homogenization of the hepatic tissue in TRIzol. Then, RNA quality was controlled with the High Sensitivity RNA Assay at the 2100 Bioanalyzer (Agilent, cat. #G2939BA) and the extracted RNA was stored at -80 °C until the RNA-seq analysis. Libraries were prepared from the extracted RNAs using the QuantSeq 3’mRNA-Seq Library Prep Kit-FWD (Cat. No. LX01596 Lexogen, Vienna, Austria) using 1 μg of RNA per library and following the manufacturers’ instructions. We modified the standard protocol by adding Unique Molecular Identifiers (UMI) during the second strand synthesis step. Indices from the Lexogen i7 6nt Index Set and i5 6nt Dual Indexing Add-on Kits (Cat. No. 044.96 and 047.96, Lexogen) were used, and 15 cycles of library amplification were performed. Libraries were eluted in 30 μL of the kit’s Elution Buffer. The double-stranded DNA concentration was quantified using the Qubit dsDNA HS Assay Kit (ThermoFisher), ranging from 3 to 12 ng/μL. The molar concentration of cDNA molecules in the individual libraries was calculated from the double-stranded DNA concentration and the single library average size (determined on a PerkinElmer Labchip GX). An equal number of cDNA molecules from each library were pooled and the final pool was purified once more in order to remove any free primer and prevent index-hopping. The pooled libraries were sequenced in a Novaseq 6000 instrument (Illumina, San Diego, CA) on an SP flowcell, producing 900M single reads 100nt.

Sequencing reads from the resulting FASTQ files were aligned onto mouse reference genome (GRCm38 primary genome assembly) using STAR aligner version 2.7.7a (Dobin et al., 2013), setting the parameters outFilterScoreMinOverLread and outFilterMatchNminOVerLread to the value 0.33. Resulting SAM files were sorted and converted to BAM files using SAM Tools (Li et al., 2009). Transcripts counts were computed using the featureCounts function available from the Rsubread R package (Liao et al., 2019), utilizing mouse gene annotation (GRCm38) for reads summarization. 6 and 12 months adult chronic rapamycin treatment RNA-seq data counts were downloaded from Gene Expression Omnibus (GEO) under accession number GSE131754 (Tyshkovskiy et al., 2019). Transcripts with a raw count lower than 20 in all biological replicates across the considered conditions were excluded. TMM (Trimmed Mean of M values) normalization and CPM (Counts Per Million) conversion were performed to obtain normalized transcript levels.

### Differential Expression Analysis

The edgeR R package (Robinson et al., 2010) was used to perform differential expression analysis. Rapamycin treated samples were compared against the respective control samples. Transcripts with a log2 fold-change higher/smaller than 1/-1, an FDR-corrected p-value <0.05 and a mean log2 CPM >0 across the replicates of at least one of the two compared groups were considered as significant differentially expressed genes (DEGs). To account for potential effects due to biological replicates generated in different sequencing batches, differential expression analysis of P4-P30 male and female mice was performed considering the batch as a covariate in the analysis model.

### Gene Set Enrichment Analysis

Gene set enrichment analysis (GSEA) of Gene Ontology (GO) Biological Processes (BP), Molecular Functions (MF) and Cellular Components (CC) terms was performed with the gseGO function of the clusterProfiler R package (Yu et al., 2012) and p-values were FDR-corrected.

As described in Tyshkovskiy et al., 2019 the input for the gseGO function was obtained by pre-ranking the list of genes of the edgeR output based on the -log10(adjusted p-value) corrected by the sign of regulation (1, -1 or 0 if the log2FC value is positive, negative or equal to 0, respectively), as:

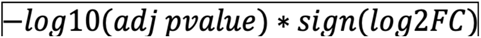

Significance scores of enriched functions were obtained from the output of the gseGO function based on -log10(q-values) corrected by the sign of the normalized enrichment score (NES), as:

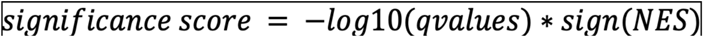

Significance scores barplots were inspected manually to choose for terms that are statistically significant in the P4-P30 Male or Female categories.

### Metabolomics on mouse livers

Metabolomic data were obtained by liquid chromatography coupled to tandem mass spectrometry. We used an API-3500 triple quadrupole mass spectrometer (AB Sciex, Framingham, MA, USA) coupled with an ExionLC™ AC System (AB Sciex). Cells were smashed in a tissue lyser for 2 min at maximum speed in 250µl of ice-cold methanol/water/acetonitrile 55:25:20 containing [U-^13^C_6_]-glucose 1ng/µl and [U-^13^C_5_]-glutamine 1ng/µl as internal standards. Lysates were spun at 15,000g for 15 min at 4°C and supernatants were then passed through a regenerated cellulose filter (4mm Ø, Sartorius). Samples were then dried under N2 flow at 40°C and resuspended in 125µl of ice-cold methanol/water 70:30 for subsequent analyses.

Amino acids, their derivatives and biogenic amine quantification was performed through previous derivatization. Briefly, 25µl out of 125µl of samples were collected and dried separately under N2 flow at 40°C. Dried samples were resuspended in 50µl of phenyl-isothiocyanate (PITC), EtOH, pyridine and water 5%:31.5%:31.5%:31.5% and then incubated for 20 min at RT, dried under N2 flow at 40°C for 90 min and finally resuspended in 100µl of 5mM ammonium acetate in MeOH/H2O 50:50. Quantification of different amino acids was performed by using a C18 column (Biocrates, Innsbruck, Austria) maintained at 50°C. The mobile phases for positive ion mode analysis were phase A: 0.2% formic acid in water and phase B: 0.2% formic acid in acetonitrile. The gradient was T0: 100%A, T5.5: 5%A, T7: 100%A with a flow rate of 500µl/min. All metabolites analyzed in the described protocols were previously validated by pure standards and internal standards were used to check instrument sensitivity.

Quantification of energy metabolites, cofactors and nucleotides was performed by using a cyano-phase LUNA column (50mm x 4.6mm, 5µm; Phenomenex) by a 5 min run in negative ion mode with two separated runs. *Protocol A*; mobile phase A was: water and phase B was: 2mM ammonium acetate in MeOH and the gradient was 10% A and 90% B for all the analysis with a flow rate of 500µl/min. *Protocol B*; mobile phase A was: water and phase B was: 2mM ammonium acetate in MeOH and the gradient was 50% A and 50% B for all the analysis with a flow rate of 500µl/min.

Acylcarnitines, GSH, GSSG and SAMe quantification was performed on the same samples by using a Varian Pursuit XRs Ultra 2.8 Diphenyl column (Agilent). Samples were analyzed by a 9 min run in positive ion mode. Mobile phases were A: 0.1% formic acid in H20 B: 0.1% formic acid in MeOH and the gradient was T0: 35%A, T2.0: 35%A, T5.0: 5%A, T5.5: 5%A, T5.51: 35%A, T9.0: 35%A with a flow rate of 300µl/min.

Bile acids were analyzed on an API-4000 triple quadrupole mass spectrometer (AB Sciex) coupled with a HPLC system (Agilent) and CTC PAL HTS autosampler (PAL System). The LC mobile phases were (A) 10mM NH_4_Ac and 0.015% formic acid in water, (B) 10mM NH_4_Ac and 0.015% formic acid in Acetonitrile/Methanol/Water (65/30/5). The gradient was as follows: T0: 65% A (flow rate 400µl/min), T0.7: 60% A (flow rate 400µl/min), T3 55% A (flow rate 400µl/min), T3.2: 45% A (flow rate 400µl/min), T5.5: 35% A (flow rate 450µl/min), T6.5: 0% A (flow rate 500µl/min), T8.5: 0% A (flow rate 600µl/min), T8.6: 65% A (flow rate 700µl/min) and T11: 65% A (flow rate 400µl/min). The Hypersil GOLD column C18 (100mm x 3 mm, 3 µm) was maintained at 50°C. The mass spectrometer was operated in negative ion mode and in selective ion monitoring SIM/SIM mode.

### MultiQuant™ software (version 3.0.3, AB Sciex) was used for data analysis and peak review of chromatograms

Raw areas were normalized by the median of all metabolite areas in the same sample. Specifically, we defined the relative metabolite abundance 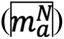 as:

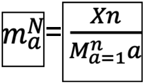

where *Xn* represents the peak area of metabolite n for samples a, b, …, z, and 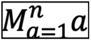 represents the median of peak areas of metabolite n for samples *a, b, …, z*. Obtained data were then transformed by log10-transformation and Pareto scaled to correct for heteroscedasticity, reduce the skewness of the data, and reduce mask effects (Ghaffari et al., 2019). In detail, obtained values were transformed by log10 and then scaled by Pareto’s method as follows:

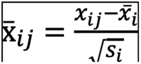

where *x_ij_* is the transformed value in the data matrix (*i* (metabolites), *j* (samples)) and *s_i_* is the standard deviation of transformed metabolite values (van den Berg et al., 2006). Obtained values were considered as relative metabolite levels. Data processing and analysis were performed by MetaboAnalyst 5.0 web tool (Chong et al., 2019).

### Frailty Index Assessment

Frailty Index assessment was performed at 7, 15 and 20 months on P4-P30 and P30-P60 rapamycin treated mice. The clinical FI score for each mouse was calculated using a modified version of the tool published previously by Whitehead et al. (Whitehead et al., 2014). Briefly, mice were placed in a fresh cage and moved to a dedicated small animal procedure room designed for behavioral testing. Mice were weighed and then a series of non-invasive observations on 30 clinical items were taken (as listed in Table S1). Clinical assessment included evaluation of the integument, musculoskeletal system, vestibulocochlear and auditory systems, ocular and nasal systems, digestive system, urogenital system, respiratory system, signs of discomfort, as well as the body weight (grams). A complete list of the clinical signs of deterioration and/or deficits evaluated in this study can be found in Table S1.

### Forelimb grip strength

A grip strength meter system (Cat. No. 47200, Ugo Basile Srl) was used to assess grip strength at 7, 15 and 20 months on P4-P30 and P30-P60 rapamycin treated mice. Mice were allowed to hold on to the grid with the front paws. Each mouse was given 3 trials over the course of 5 minutes. Average values (gF) were then normalized using the weight (g) to calculate the grip strength of an individual mouse.

### Drosophila stocks

*Drosophila* strains were raised on standard cornmeal and molasses medium in fly food vials (25 x 95 mm, Biosigma, cat. #789008) at 25°C where not otherwise indicated. Lyophilised Nutri-Fly Food (Genesee Scientific, cat. #789211), was used during the rapamycin treatment window. The stocks used in this study were: *white iso31 (w^iso31^* in the text, kind gift from Alex Gould), *y[1] w[*]; P{w[+mC]=UAS-mCD8::GFP.L}LL5, P{UAS-mCD8::GFP.L}2* (BDSC_5137 - *UAS-GFP* in the text), *P{w[+mC]=tubP-GAL80[ts]}10; P{w[+mC]=tubP-GAL4}LL7/TM6B, Tb[1]* (BDSC_86328 - *TubGal80;TubGal4* in the text), *y[1] w[*]; P{w[+mC]=tubP-GAL4}LL7/TM3, Sb[1] Ser[1]* (BDSC_5138 - TubGal4 in the text) purchased from Bloomington Drosophila Stock Center and *M{UAS-St1.ORF.3xHA.GW}ZH-86Fb* (cat. #F003139, *UAS-dST1* in the text) purchased from FlyORF.

### Rapamycin treatment and survival analysis in *Drosophila*

#### Larval transient treatment

Rapamycin (Alfa Aesar, cat. #J62473) was dissolved in ethanol at a final concentration of 20 mg/mL and then working aliquots were diluted in milli-Q water up to a concentration of 1 μM, 50 μM and 200 μM. For each experiment of survival analysis, three independent cohorts of *w^iso31^* flies were established at 25 °C on Nutri-Fly food, to favor the distribution of rapamycin within the food. Adult flies were allowed to lay eggs in the test tubes for 12 hours leading to egg-synchronization. 72 hours after egg-laying, 1 ml of 1 μM, 50 μM or 200 μM rapamycin solutions or ethanol as control was added drop by drop in each tube. Newly eclosed flies where promptly collected, divided by gender and maintained in new vials containing standard food at a density of 10 ± 3 flies per vial. Flies were flipped to new vials at least twice a week and monitored for death events.

#### Adult early transient treatment

Five cohorts of *w^iso31^* flies were established on standard food. Adult flies were allowed to lay eggs in the test tubes for 12 hours leading egg-synchronization. Newly eclosed flies where collected, divided by gender and maintained in new vials containing standard food at a density of 10 ± 3 flies per vial. **From day 0 to 10**: On the first day post-eclosion, flies were transferred to new vials containing Nutri-Fly food supplemented with 1 ml of ethanol or 200 μM of rapamycin. Flies were maintained in the Nutri-Fly food for 10 days (flipped three times) and then transferred back to fresh vials with standard food. **From day 10 to 13**: On the tenth day post-eclosion, flies were transferred to new vials containing Nutri-Fly food supplemented with 1 ml of ethanol or 200 μM of rapamycin. Flies were maintained in the Nutri-Fly food for 3 days and then transferred back to fresh vials with standard food. **From day 10 to 20**: On the tenth day post-eclosion, flies were transferred to new vials containing Nutri-Fly food supplemented with 1 ml of ethanol or 200 μM of rapamycin. Flies were maintained in the Nutri-Fly food for 10 days (flipped three times a week) and then transferred back to fresh vials with standard food. For all the timepoints listed above flies were flipped to new vials at least twice a week and monitored for death events.

#### *Drosophila* ‘s food receipts

Flies were reared on standard sugar/yeast/agar food (Standard food) or Nutri-Fly food, based on the experimental setup. Standard food consists of 85 g corn flour, 60 g sugar, 30g Brewers Yeast (ACROS Organics™, cat. # 368080050), 10g Insectagar ZN5 (B.&V. S.R.L. The Agar Company), 50g Molasses, 15 ml/L 1M Propionic Acid (Genesee Scientific, cat. #789177), 15 ml/L 10 % w/v Tegosept in Et-OH 96 % (Genesee Scientific, cat. #789063) per liter. Lyophilised food consists of 200 g Nutri-Fly Food (Genesee Scientific, cat. #789211), 16,25 g Brewers Yeast (ACROS Organics™, cat. # 368080050) per liter.

### Transient overexpression of dST1

#### Larval transient overexpression

To study the effect of transient dST1 overexpression on *Drosophila* lifespan, the following crosses were established on standard food: *;TubulinGal80^ts^;TubulinGal4/TM6B* flies with *;;UAS- dST1 flies*, and *;TubulinGal80^ts^;TubulinGal4/TM6B* flies with *;UAS-GFP/Cyo;* flies. Adult flies, maintained at 18°C, were allowed to lay eggs in the test tubes for 12 hours leading to egg-synchronization. 130 hours after egg-laying, vials containing developing larvae were shifted to 29 °C, to induce the temperature-sensitive inhibition of the Gal80 and allow the overexpression of the genes of interest (dST1 or GFP) till puparium formation and then maintained at 18 °C till death. Adult flies of each cross were selected for the following genotypes: *;TubulinGal80^ts^/+;TubulinGal4/UAS-dST1* (in the text *TubGal80;TubGal4;UAS-dST1*) and *;TubulinGal80^ts^/UAS-GFP;TubulinGal4/+* (in the text *TubGal80;TubGal4;UAS-GFP*). Selected flies were divided by gender in new vials containing standard food, at a density of 10 ± 3 flies per vial. Flies were flipped at least twice a week and monitored for death events.

### Constitutive dST1 overexpression

To study the effect of constitutive dST1 overexpression on *Drosophila* lifespan, the following crosses were established on standard food: *;;TubGal4/TM6B flies with ;;UASdST1 flies, ;;TubGal4/TM6B flies with ;UAS-GFP/Cyo;* flies. Adults were allowed to lay eggs in the test tubes for 12 hours leading to egg-synchronization. Male and female flies were selected after eclosion based on the specific genotype for the cross (*;;TubGal4;UAS-dST1*, *;;TubGal4;UAS-GFP*,and maintained at 25°C in new vials. Flies were flipped three times a week and monitored for death events.

### Antibodies

Primary antibodies used are listed here: Rabbit anti-Phospho-S6 Ribosomal Protein (Ser235/236) (1:1000, Cell Signaling, cat. #2211), Rabbit S6 Ribosomal Protein (5G10) (1:4000, Cell Signaling, cat. #2217). anti-dS6K (generous gift from Aurelio Teleman 1:3,000), and anti-phospho-Thr398-S6K (Cell Signaling Technologies #9209, 1:1,000), The secondary antibody used is Peroxidase AffiniPure Goat Anti-Rabbit IgG (H + L) (1:5000, Jackson ImmunoResearch, cat. #111-035-003) and Rabbit anti-Guinea Pig IgG (H+L) Secondary Antibody (1:5000 thermo fisher cat.# HRP 61-4620).

### RNA Isolation and Real-Time PCR Analysis

*Drosophila*: Three biological replicates of L3 wondering larvae (n=30) and 10-days-old flies (n=10) treated for 12 hours with rapamycin or ethanol were snap frozen and stored at -80 °C. Total RNA from frozen samples was isolated with TRIzol Reagent (Invitrogen), using pellet pestles (Sigma-Aldrich, cat. #Z359971-1EA). RNA was reverse transcribed using iScript cDNA synthesis kit (Biorad, cat. #1708891) according to the manufacturer’s instructions and quantitative PCR was performed using Power SYBR Green PCR Master Mix (Applied Biosystems, cat. #4367659). The Ct values were normalized to the housekeeping gene *tubulin*. Primer sequences used for qRT-PCR are listed in Table S7.

### Climbing performance assay

Adult flies were placed in a plastic cylinder. Cylinder was tapped quickly and it was recorded the number of flies over 5 centimeters after 15 seconds. This step was repeated three times.

### Statistical analysis

Survival analysis was performed calculating the lifespan in days for every mouse or *Drosophila* and data were displayed using the Kaplan-Meier curve. The statistical significance of the results was tested using the Log-rank (Mantel-Cox) test.

Data of weight and length analyses are presented as mean + SD of three litters. Two-tailed Student’s t-test was used for calculating significance values between treated and control mice.

For qRT-PCR analysis, data are presented as mean + s.e.m. of three biologically independent samples and a two-tailed Student’s t-test was used to calculate significance.

For forelimb grip strength and FI analyses, data are presented as mean + s.e.m. and a two-way anova was used to calculate significance.

## List of Tables

Table S1 Table S1. Summary table of Frailty index analysis at 7, 15 and 20 months for P4-P30 and P30-P60 rapamycin treated mice, related to Figure 2.

Table S2 Differentially expressed genes (DEGs) after rapamycin treatment across the different time-windows processed at the last day of treatment, related to Figures 3, 4 and figure S2.

Table S3 Differentially expressed genes (DEGs) after rapamycin treatment across the different time-windows processed at P350, related to Figures 3.

Table S4 Significant GO terms result of GSEA of rapamycin treated mice across the different time-windows processed at the last day of treatment, related to Figure 4 and Figure S2.

Table S5 Metabolite profiling data of rapamycin treated mice across the different time-windows and timepoints. Related to Figure 5 and fig. S3

Table S6 Summary table of *Drosophila melanogaster* experiments related to Figure 6,7 and figure S4

Table S7 qRT-PCR primers related to figures S6.

## Figure Legends

**Figure S1.**
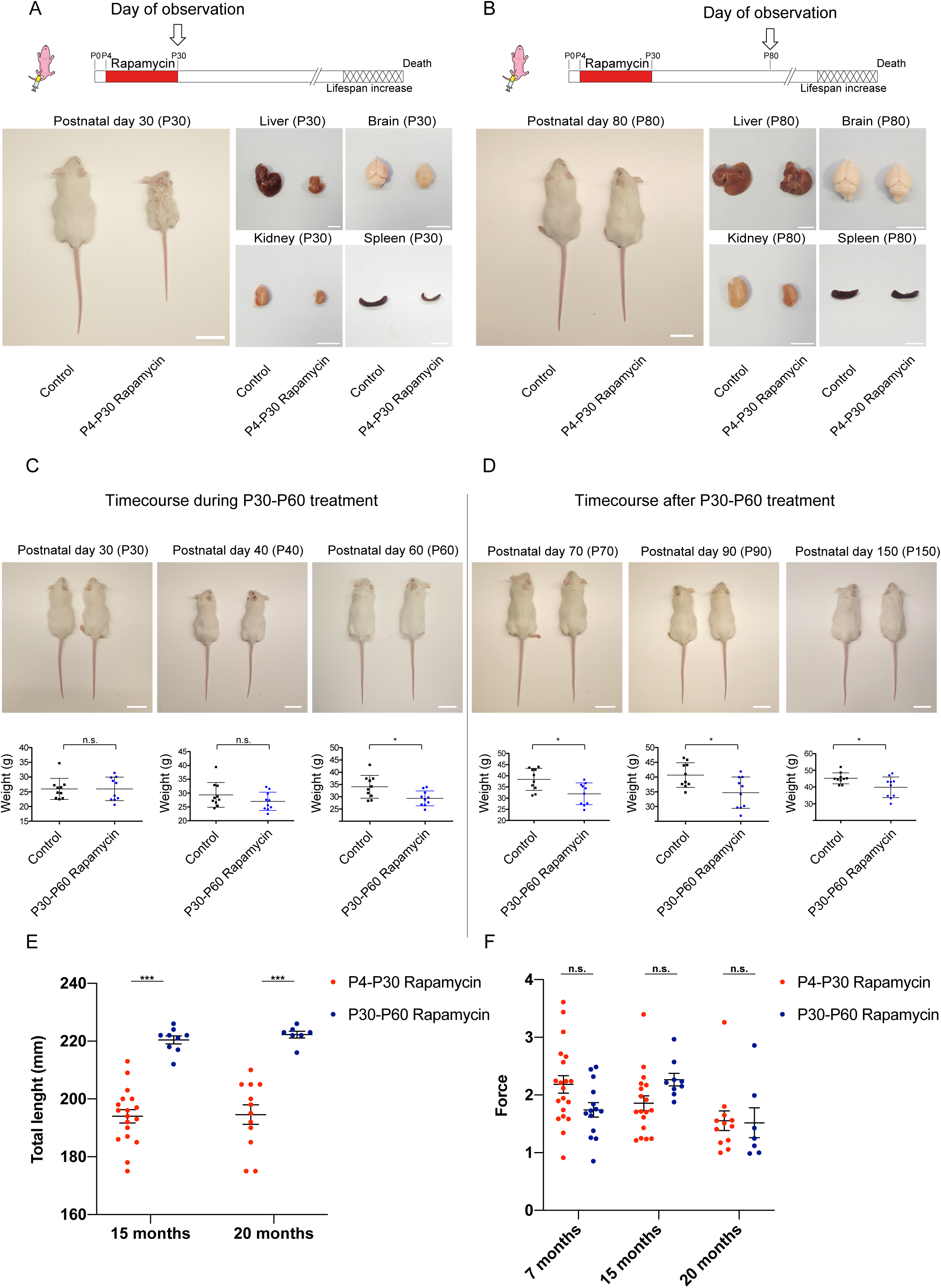
Analysis of physical status of P4-P30 and P30-P60 rapamycin treated mice at different timepoints. **(A, B)** Upper panels: Schematic illustration of the experimental procedures. Rapamycin effects on body and organs size in P4-P30 treated mice. Representative images of liver, brain, kidney and spleen have been taken at P30 (A) and P80 (B). Scale bar: 3 cm **(C)** Upper panels: Representative images showing rapamycin effects on mice body size during P30-P60 treatment, taken at P30, P40 and P60 (end of treatment). Scale bar: 3 cm. Lower panels: Scatter dot plot indicating body weight of control and P30-P60 rapamycin treated mice during treatment. Data are indicated as mean + SD. **(D)** Upper panels: Representative images showing rapamycin effects on mice body size after P30-P60 treatment, taken at P70, P90 and P150. Scale bar: 3 cm. Lower panels: Scatter dot plot indicating body weight of control and P30-P60 rapamycin treated mice after treatment. Data are indicated as mean + SD. **(E)** Box plots indicating the total length (mm) of P4-P30 and P30-P60 rapamycin treated mice. Red scatter dots indicate the measurements of P4-P30 rapamycin mice at 15 months (n=18), and 20 months (n=12). Blue scatter dots indicate the measurements of P30-P60 rapamycin mice at 15 months (n=9) and 20 months (n=7). Data are indicated as mean + SEM. **(F)** Box plots indicating the force (Newton/grams) resulted from grip strength analysis of P4-P30 and P30-P60 rapamycin treated mice. Red scatter dots indicate the measurements from P4-P30 rapamycin mice at 7 months (n=20), 15 months (n=18) and 20 months (n=12). Blue scatter dots indicate the measurements from P30-P60 rapamycin mice at 7 months (n=14), 15 months (n=9) and 20 months (n=7). Data are indicated as mean + SEM. *p-value<0.05, **p-value<0.005, ***p-value<0.0005, n.s.-not significant.

**Figure S2.**
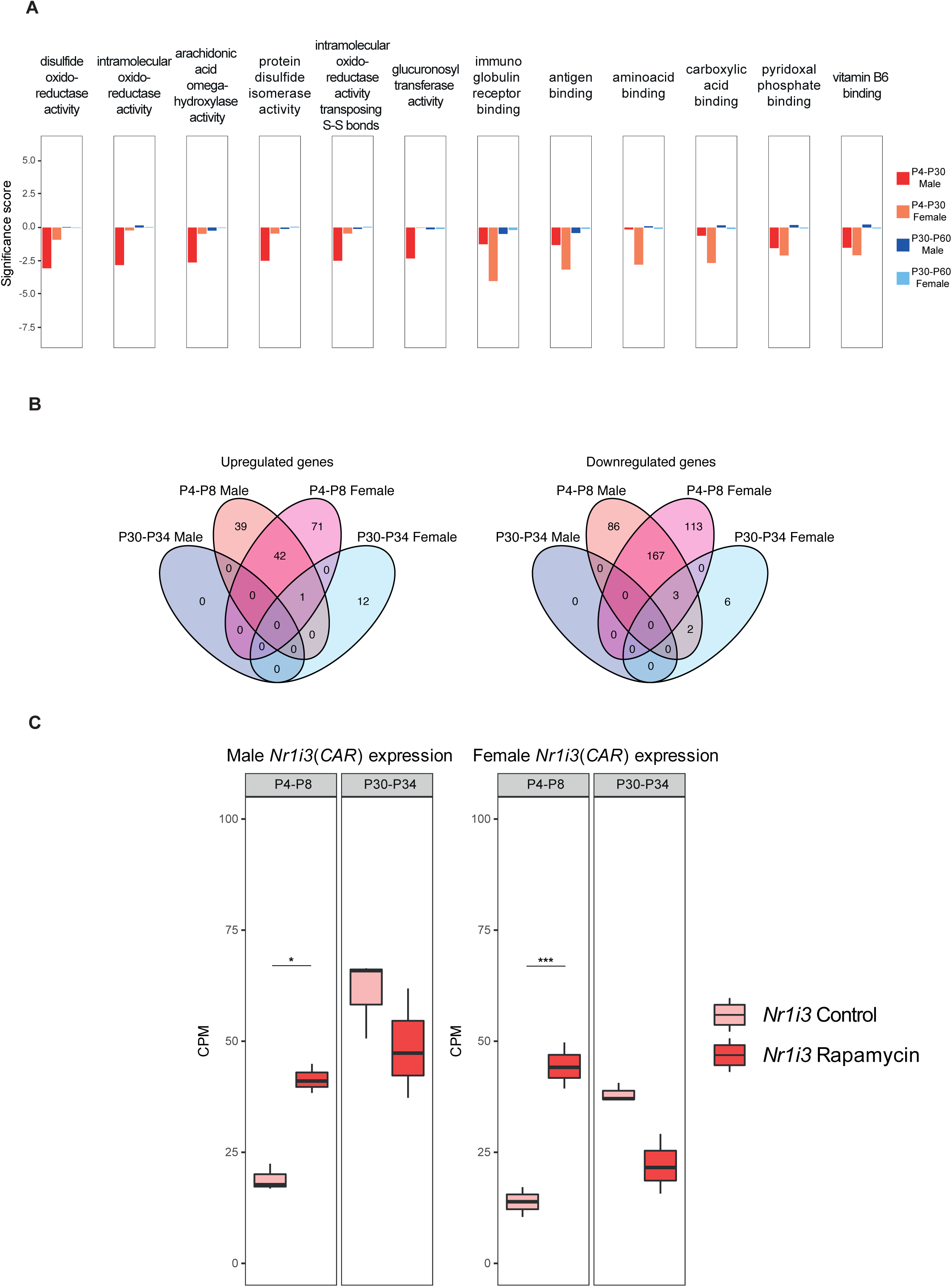
RNA-seq analysis of acute rapamycin treatments (P4-P8 and P30-P34). **(A)** Gene set enrichment analysis (GSEA) results of P4-P30 and P30-P60 treatments in male and female mice. Significance score, calculated as log10(q-value) corrected by the sign of regulation, is plotted on the y axis. The whole list of enriched GO terms is available in Table S4. **(B)** Landscape of up- and down-regulated genes across the P4-P8 and P30-P34 treatments in male and female mice. Venn diagrams are used to highlight private and shared differentially expressed genes. **(C)** Expression profile of *Nr1i3* (*CAR*) gene across P4-P8 and P30-P34 male (left side) and female (right side) treated mice. Values in treated and control samples across the different conditions are shown as median CPM with bars representing standard deviations across the biological replicates. *p-value<0.05, **p-value<0.005, ***p-value<0.0005, n.s.-not significant.

**Figure S3.**
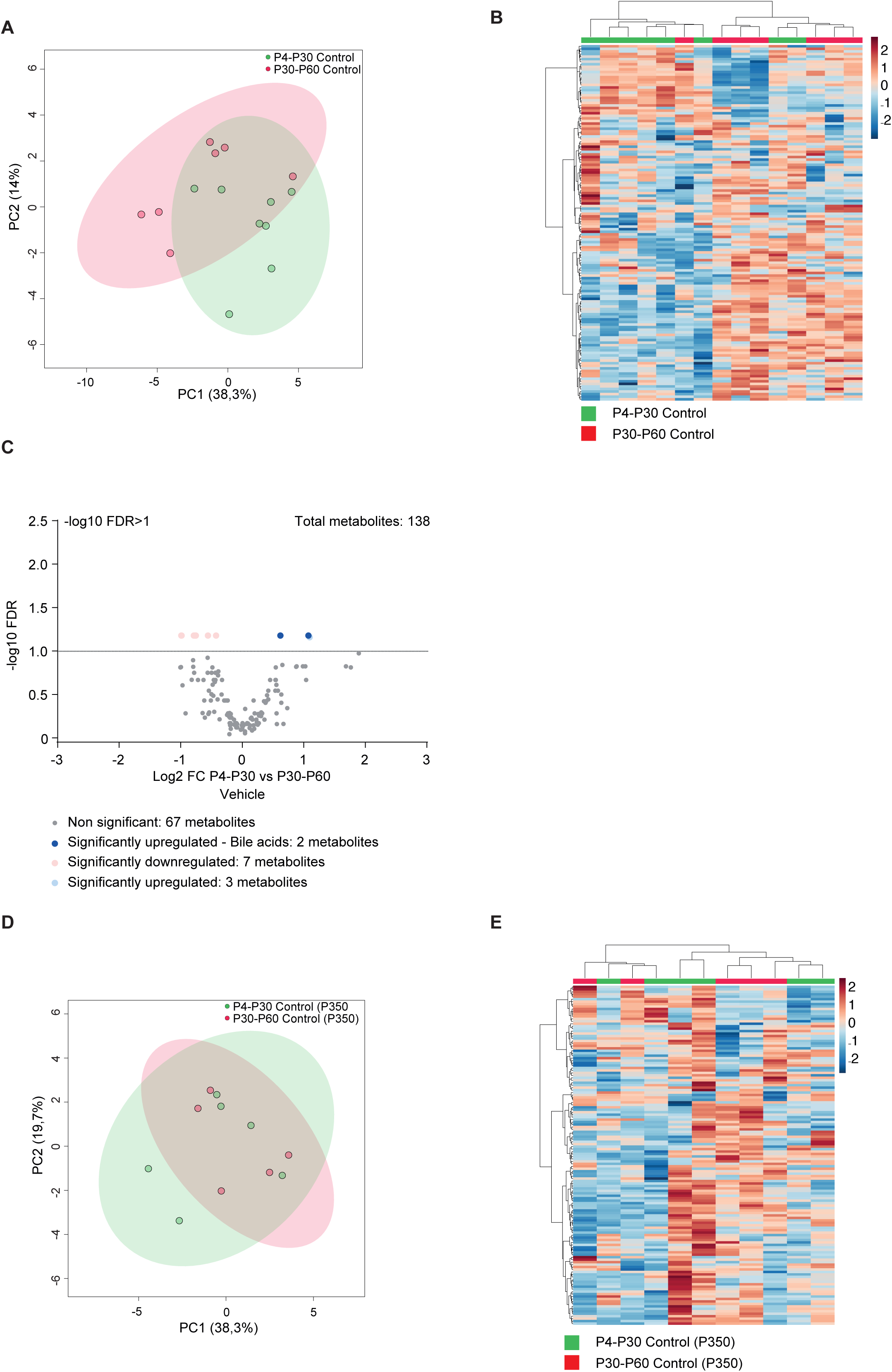
Metabolic changes in early transient treated control mice. **(A, B)** Principal Component Analysis (PCA) (**A**) and heatmap (**B**) of liver metabolomic profile from P4-P30 (green samples) and P30-P60 (red samples) mice treated with vehicle. **(C)** Volcano plots showing the metabolomic changes in P4-P30 compared to P30-P60 mice treated with vehicle. Each circle represents one metabolite. The log2 fold change is represented on the x-axis. The y-axis shows the -log10 of the FDR. A FDR of 0.1 is indicated by gray line. Grey, pink and light blue dots represent unchanged, significantly downregulated and significantly upregulated metabolites, respectively. Blue dots represent significantly upregulated bile acids in P4-P30 compared to P30-P60 mice treated with vehicle. **(D, E)** Principal Component Analysis (PCA) (**D**) and heatmap (**E**) of liver metabolomic profile from P4-P30 (green samples) and P30-P60 (red samples) mice transiently treated with vehicle and analyzed at P350.

**Figure S4.**
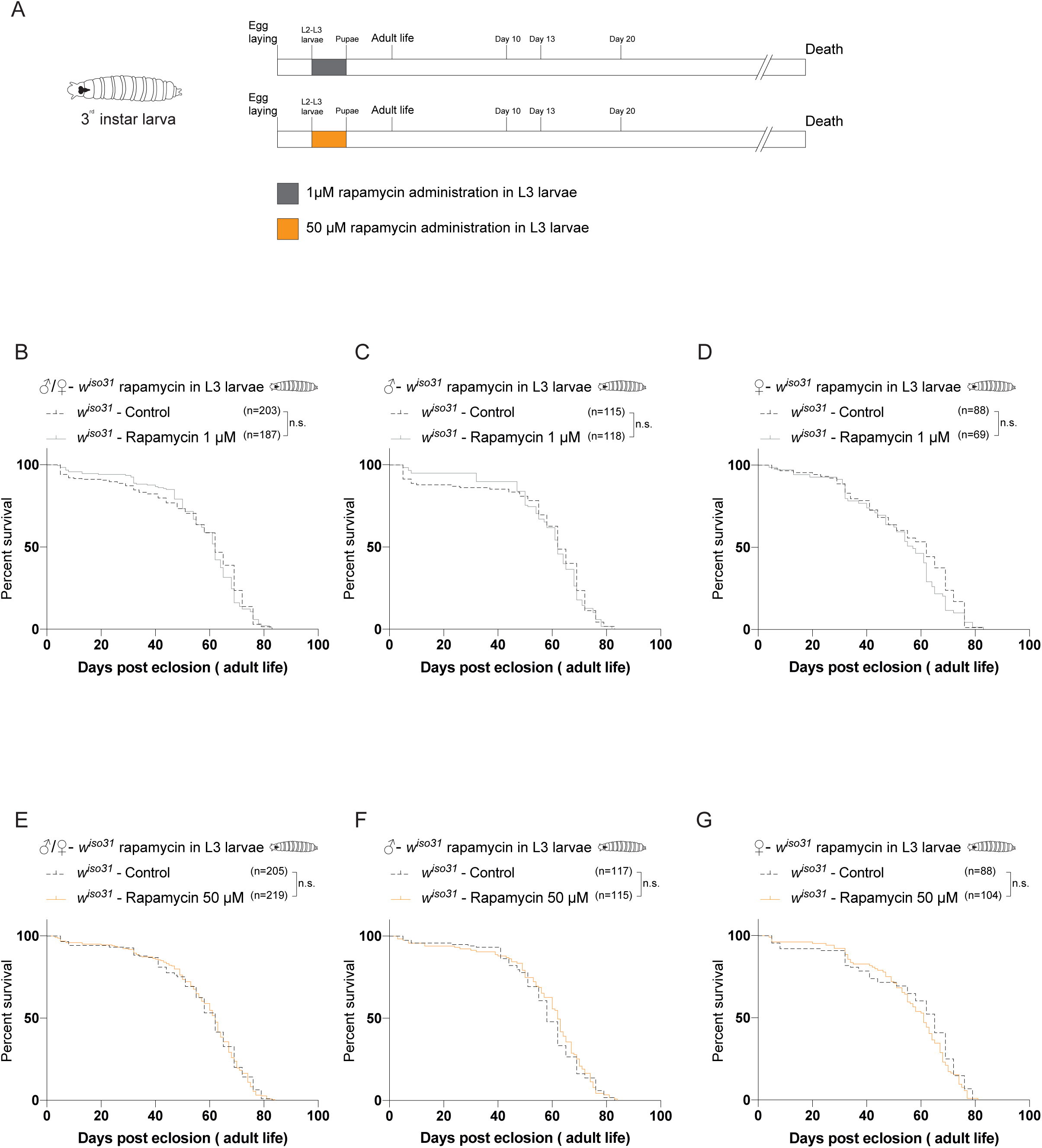
Early transient 1 µM and 50 µM rapamycin treatment does not induces a time-dependent effect on lifespan. **(A)** Schematic illustration of the experimental procedure and results. Flies were transiently treated during larval stages from 72 hours after egg-laying to puparium formation with 1 µM (grey) or 50 µM rapamycin (yellow). **(B)** Survival curves of *w^iso31^* flies transiently treated from 72 hours after egg-laying till puparium formation (males + females) with Et-OH (control) or rapamycin 1 µM. **(C, D)** Survival curves of male (**C**) and female (**D**) *w^iso31^*flies transiently treated from 72 hours after egg-laying till puparium formation with Et-OH (control) or rapamycin 1 µM. **(E)** Survival curves of *w^iso31^* flies transiently treated from 72 hours after egg-laying till puparium formation (males + females) with Et-OH (control) or rapamycin 50 µM. **(F, G)** Survival curves of male (**F**) and female (**G**) *w^iso31^*flies transiently treated from 72 hours after egg-laying till puparium formation with Et-OH (control) or rapamycin 50 µM. *p-value<0.05, **p-value<0.005, ***p-value<0.0005, n.s.- not significant.

**Figure S5.**
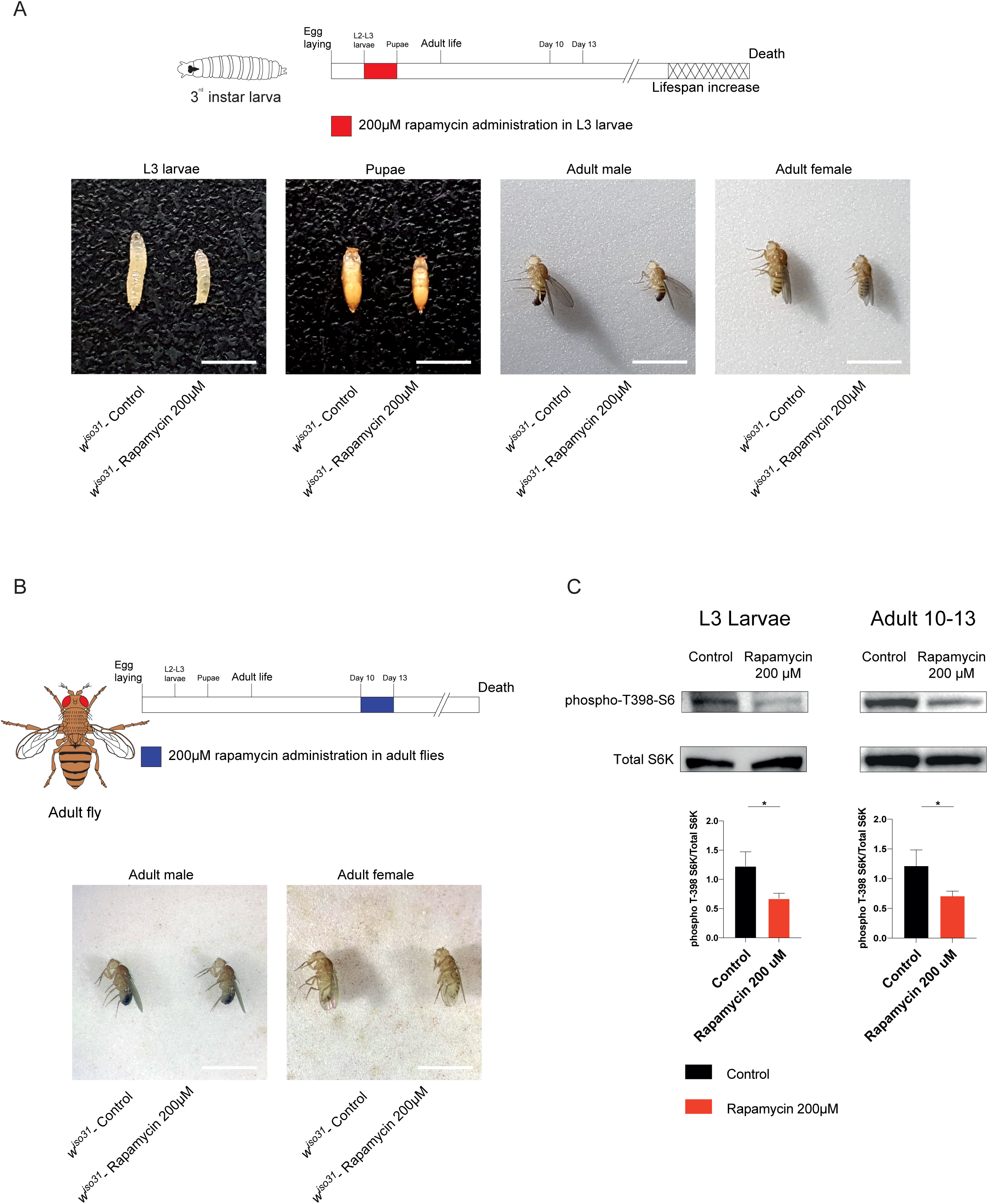
Physical status and biochemical effectiveness of rapamycin treatment in *w^iso31^ Drosophila melanogaster*. **(A)** Representative images showing rapamycin effects on *Drosophila* body size during and after treatment on third instar (L3) larvae. Images acquired during L3 larvae (120 hours after egg-laying), pupae and adult (1 day post eclosion) stages. Scale bar: 3 mm **(B)** Representative images showing rapamycin effects on *Drosophila* body size after treatment on 10-days-old flies. Images acquired during adult stage (3 days post treatment). Scale bar: 3 mm **(C)** Western blot analysis of S6 Ribosomal Protein and phospho-T398-S6 Ribosomal Protein on whole-flies protein extracts of L3 Larvae (left side) and 10-days-old flies treated for 3 days (right side). *p value<0.05.

**Figure S6.**
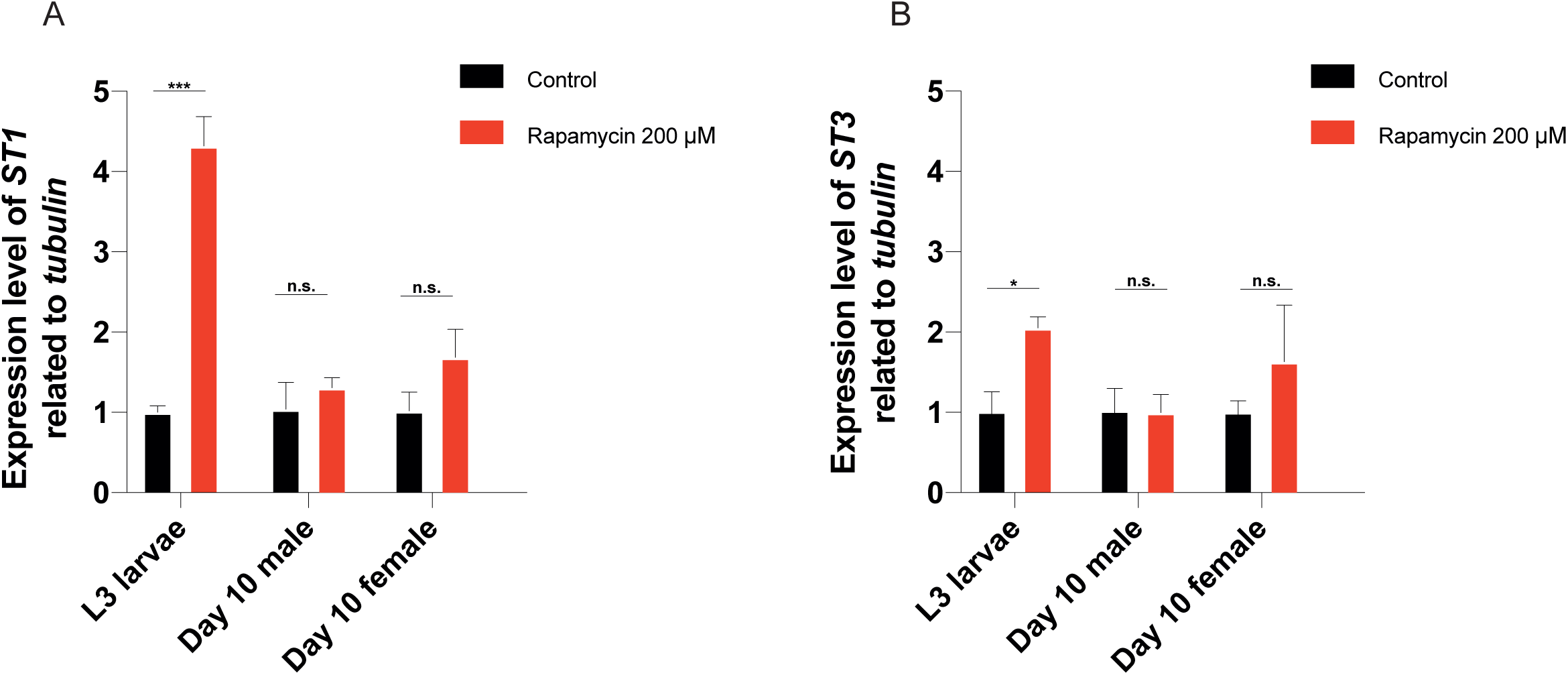
Sulfotransferases are putative rapamycin targets during larval stages but not in adult life. **(A, B)** Gene expression analysis via qRT-PCR of *dST1***(A)** and *dST3***(B)** in *w^iso31^* L3 larvae and 10-days-old adult flies treated with 200 μM rapamycin, 12 hours after treatment. Data are indicated as mean + SEM.*p-value<0.05, **p-value<0.005, ***p-value<0.0005, n.s.-not significant.

## Notes

### Competing Interest Statement

The authors have declared no competing interest.

